# Systematic empirical evaluation of individual base editing targets: validating therapeutic targets in *USH2A* and comparison of methods

**DOI:** 10.1101/2024.10.02.615422

**Authors:** Yuki Tachida, Kannan V. Manian, Rossano Butcher, Jonathan M. Levy, Nachiket Pendse, Erin Hennessey, David R. Liu, Eric A. Pierce, Qin Liu, Jason Comander

## Abstract

Base editing shows promise for the correction of human mutations at a higher efficiency than other repair methods and is especially attractive for mutations in large genes that are not amenable to gene augmentation therapy. Here, we demonstrate a comprehensive workflow for in vitro screening of potential therapeutic base editing targets for the *USH2A* gene and empirically validate the efficiency of adenine and cytosine base editor/guide combinations for correcting 35 *USH2A* mutations. Editing efficiency and bystander edits are compared between different target templates (plasmids versus transgenes) and assays (Next generation sequencing versus Sanger), as well as comparisons between unbiased empirical results and computational predictions. Based on these observations, practical assay recommendations are discussed. Finally, a humanized knock-in mouse model was created with the best-performing target, the nonsense mutation c.11864G>A p.(Trp3955*). Split-intein AAV9 delivery of editing reagents resulted in the restoration of USH2A protein and a correction rate of 65 ± 3% at the mutant base pair and of 52 ± 3% excluding bystander amino acid changes. This efficiency compares favorably to a prior genome editing strategy tested in the retina that completed a clinical trial and demonstrates the effectiveness of this overall strategy to identify and test base editing reagents with the potential for human therapeutic applications.

## Introduction

Re-writing DNA using programmable genome editing tools based on CRISPR/Cas9 technology can be leveraged to treat human genetic diseases.^1^ Conventional CRISPR/Cas9-based genome editing can address mutations where deletions or indels are therapeutic. However, the technology requires DNA double-strand breaks (DSBs) as part of the editing mechanism, which can result in genotoxic lesions, such as unwanted insertions, deletions, and vector integrations.^2,3^ Base editors (BEs) are a newer category of genome editors known for their high efficiency and can be targeted to repair single nucleotide variations (SNVs) with minimal creation of DNA double-strand breaks and indels.^4–6^ Currently, three DNA base editors have been developed, ABE (A-T:G-C conversion), CBE (C-G: T-A conversion), and GBE (C-G: G-C conversion).^7,8^ These BEs could be used to target 63% of pathogenic SNVs reported in ClinVar.^7–9^ Prime editing is a third genome editing strategy, which has more flexibility in the types of replacements that can be made, though initially it did not have as high an efficiency.^10,11^ All of these approaches are being pursued, including for inherited retinal disorders (IRDs), where effective treatment options are still lacking.

Developments in gene therapy for IRDs serve as an informative example regarding how BEs can fit into current gene therapy approaches. Initially, gene augmentation was used as a therapeutic approach for several monogenic IRDs caused by mutation of smaller genes whose coding sequences fits into adeno-associated virus (AAV) vectors.^12–15^ In fact, the first FDA-approved gene therapy product for any inherited disease, voretigene neparvovec, was approved to treat the retinal dystrophy caused by mutations in *RPE65* gene.^13,16^ This drug uses adeno-associated virus (AAV2) to deliver a full-length copy of the *RPE65* gene into the retinal pigment epithelial (RPE) cells of patients.^17,18^ Attempts to extend this approach for larger genes using the increased cargo capacity of lentivirus was not successful (NCT01505062, NCT02065011),^19^ possibly attributable to low transduction rates.

An alternative approach is to edit the DNA of large genes in situ, e.g., in the target cells’ genomes. *CEP290*, a larger gene, was one of the first to reach human clinical trials for in vivo genome editing. Sub-retinal injection of AAV5 carrying SaCas9 and two guide RNAs against a *CEP290* cryptic splice site resulted in a correction rate that met the therapeutic threshold ex-vivo;^20^ clinical trial results recently showed a benefit in some patients.^1^ However, this approach is only applicable to mutations that can be addressed by deletions. In recent years, two studies reported using lentiviral vectors to deliver ABE to correct *RPE65* mutation in the retinal pigment epithelium of a mouse model of Leber congenital amaurosis (LCA).^21,22^ However, lentiviruses are not the ideal vector for therapeutic delivery in the human retina. In additional studies, AAV was to deliver BE to the retina or retinal pigment epithelium in additional models of retinal degeneration, including rd10 (*PDE6B*) mice^23^ and *RPE65* mice^21,24^. Since gene augmentation therapy has been pursued for those two genes, the concept of base editing as a therapeutic solution for the treatment of inherited retinal diseases in humans might have the largest advantage when applied to large genes.^13,18,25,26^

This study was designed to demonstrate that the adenine and cytosine base editors (ABEs and CBEs) can be efficiently tested and applied to correct mutations in a large disease gene, *USH2A*. Mutations in *USH2A* cause retinitis pigmentosa, and in cases where hearing loss is also present, the syndrome is termed Usher syndrome type 2 (USH2).^27,28^ Pathogenic mutations in *USH2A* can lead to the degeneration of photoreceptor and developmental defects of cochlear hair cells, leading to vision loss due to retinitis pigmentosa (RP) and hearing loss due to impaired auditory and vestibular function.^29,30^ *USH2A* spans ∼800 kb in the genome and contains 72 exons, with a large coding sequence of 15.6 kb.^31,32^ Therefore, the gene augmentation approach is a substantial challenge with the available vectors, even with split-vector approaches.

The BE approach, in its simplest form, is mutation-specific. Some *USH2A* mutations are more common than others, and there are over 1,700 variants classified as pathogenic or likely pathogenic in the LOVD and HGMD databases.^33^ With the ultimate goal of correcting *USH2A* mutations in the photoreceptors and cochlear hair cells of patients with *USH2A* mutations, we hypothesize that BE using ABEs and CBEs could improve function and/or delay disease progression.

This study bioinformatically identifies candidate mutations for *USH2A* BE and empirically evaluates the efficiency in vitro and in a mouse model of *USH2A*. Following the initial discovery of BEs, several variants of editors were generated to enhance precision and efficiency.^34–36^ For editing point mutations associated with human diseases such as sickle cell disease, thalassemia, and familial hypercholesterolemia, BEs have been demonstrated to efficiently edit and produce therapeutically desired outcomes.^37–40^ The efficiency of BE depends not only on the distance of the target mutation from the protospacer adjacent motif (PAM) site,^4,41^ but also on the sequence of the sgRNA itself. Empirical testing of sgRNAs helps determine which mutations are most suitable as treatment targets using base editing. Other potential limitations include bystander editing, where one or more bases other than the target base are edited within the protospacer region of the activity window.^42^ Currently there are a broad spectrum of Cas9 variants with varying PAM specificities that may help to minimize constraints related to PAM availability and bystander editing.^42,43^ For translational purposes, the split-intein system can be used to package genes in two or more pieces, which are then spliced together at the protein level inside the target cell.^44^ We previously demonstrated the successful delivery of ABEs into the retina and other tissues using AAV split-intein constructs, achieving in vivo base editing of the *DNMT1* test locus with a therapeutically-relevant efficiency.^45^

This study performs a systematic evaluation using in vitro approaches to empirically test the editing outcome for 35 editable mutations in *USH2A* gene using a variety ABEs and CBEs editors and gRNA combinations. Notably, patient-derived cell lines carrying these mutations are not readily available. Therefore, to facilitate the testing of editing efficiencies, we established an evaluation system that compared plasmid-based and transgene-based editing. The transgene was designed as a stably integrated tandem array of 35 target site sequences. Additionally, comparisons of editing efficiency were also made between empirical results and bioinformatic predictions that were released only after the guides and editors had been selected for this study. Finally, a mouse model was created for the editor-guide combination that was the best performing and the most frequently reported among the mutations tested, to demonstrate the successful restoration of *USH2A* gene expression in vivo.

## Results

### Selection of *USH2A* mutations as candidates for BE treatment

To identify potential ABE and CBE targetable *USH2A* mutations, we compiled all previously reported *USH2A* mutations using information from the LOVD and HMGD databases.^46,47^ We narrowed this list to those that were pathogenic or likely pathogenic, and to those that have been reported more than once in the LOVD database. Out of the 1,260 mutations in *USH2A* extracted from these databases, we narrowed the list to 389 mutations that were both pathogenic and reported more than once in the LOVD database (**Figure 1A**). Then, the sequence context of each mutation was analyzed for the presence of a PAM sequence within a certain editing window. At the time of the initiation of this project, existing base editors were limited to making transition mutations: C to T, T to C, A to G, or G to A. The PAM variants included were NGG (SpCas9-WT), NGA (SpCas9-VRQR), NG (SpCas9-NG), NNGRRT (SaCas9-WT), and NNNRRT (SaCas9-KKH). Targets were also limited by the presence of a PAM sequence within a certain editing window location. The editing window was set from position 2 to 13 (-19 bp to -8 bases upstream of the 5’ end of the PAM sequence) (**Figure 1B**). 142 mutations met these constraints. We chose 35 of these mutations for empirical cell-based experiments (**Table 1**), emphasizing those targets within the most active portion of the editing window and those that had been more frequently reported in humans.

**Figure 1.**
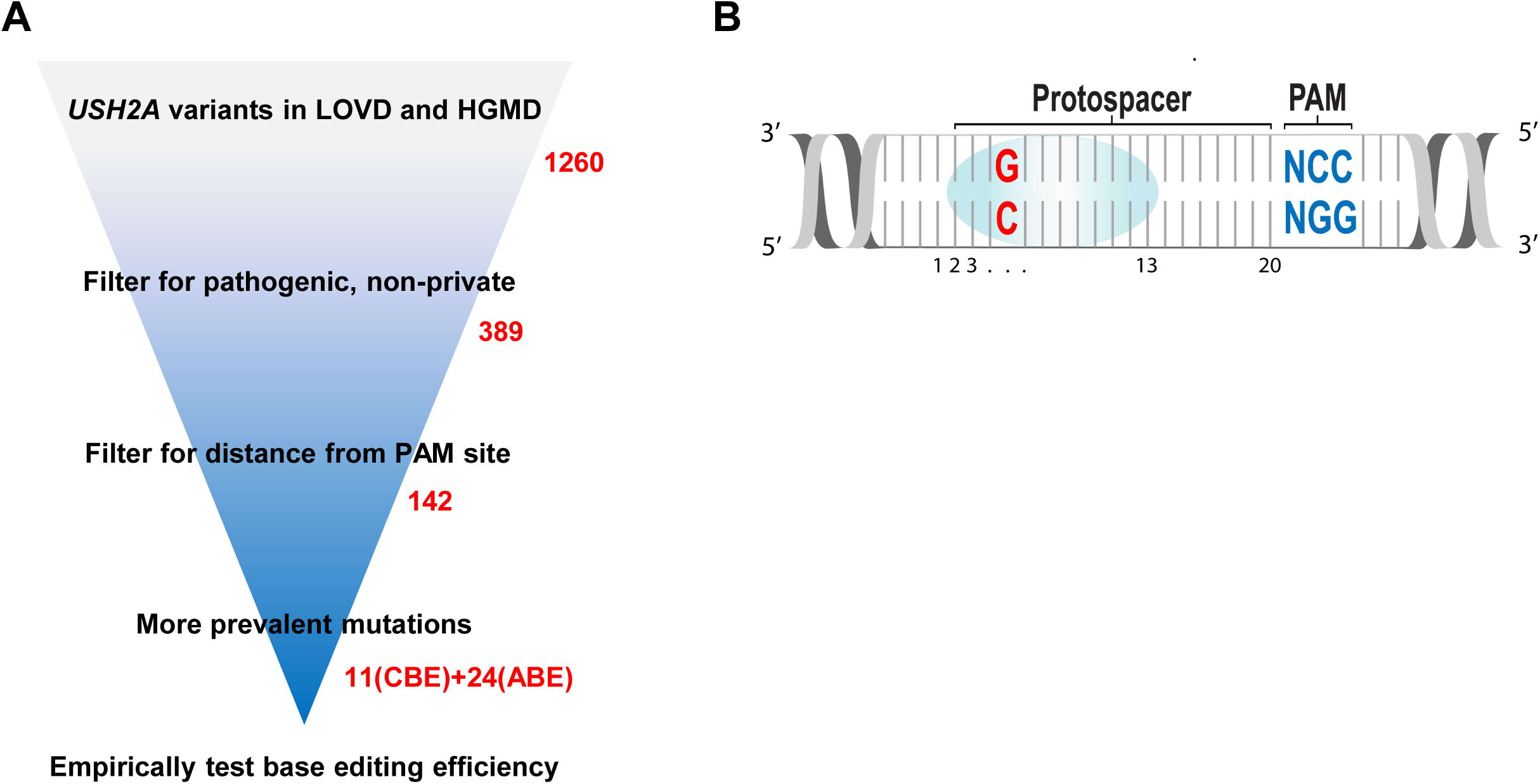
**(A)** Schema describing the *in silico* filtering of candidate USH2A mutations from LOVD and HGMD to be tested for correction by base editing. **(B)** Numbering convention used for describing base editing target locations. The bottom strand runs 5’ to 3’ and contains the target A or C to be edited and the PAM sequence.

**Table 1.**
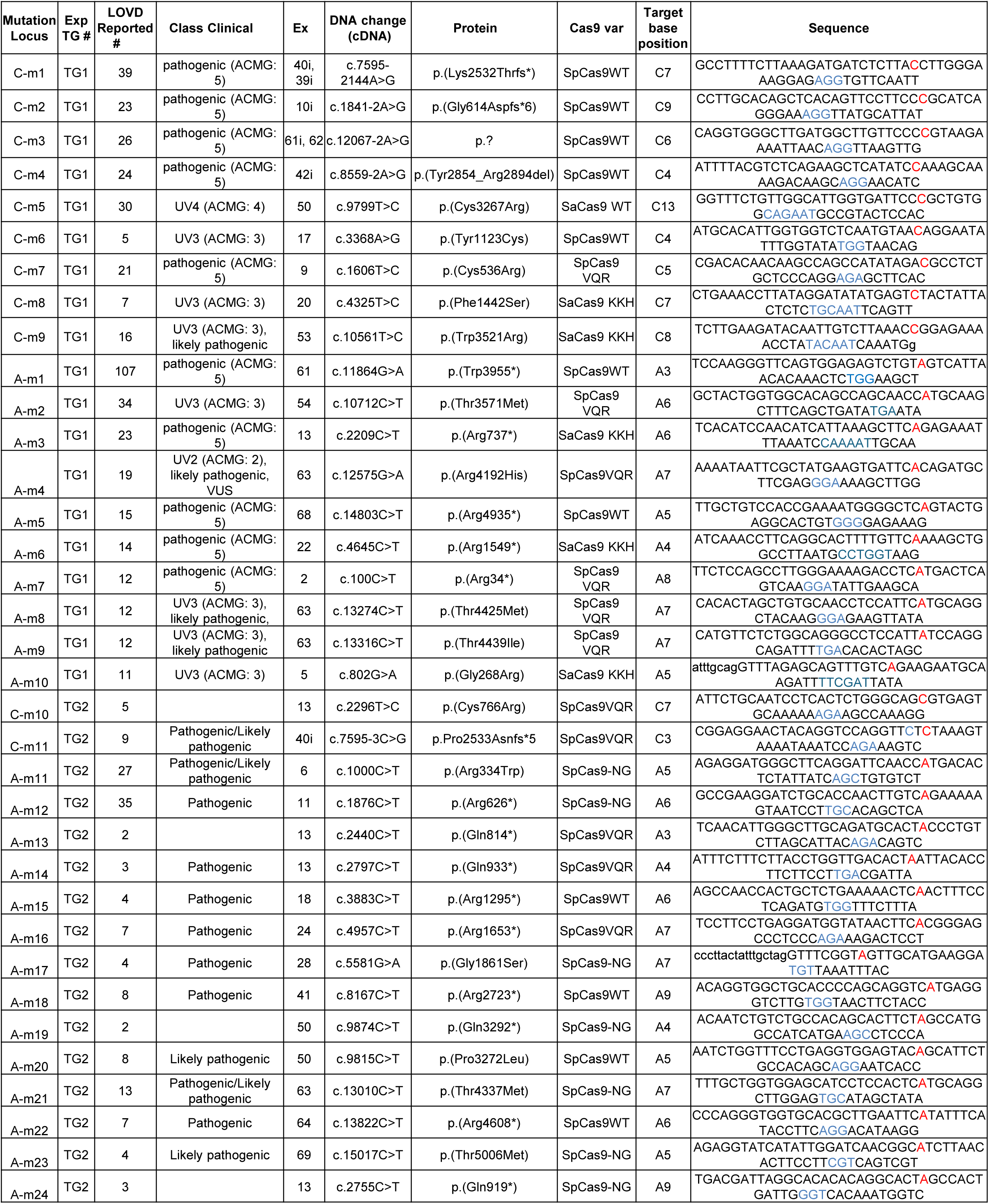
Characteristics of selected *USH2A* target mutations for base editing. The 51bp sequences of each target site are shown. The mutant base pair is shown in red, and the PAM sequence is blue.

### Generation of mutant *USH2A* transgene stable cell lines

Cell lines containing the selected target sequences are not readily available. To introduce multiple mutant sequences into cells efficiently, two synthetic constructs were designed that were composed of tandem arrays of 51 bp fragments, each containing a target mutation and adjacent sequence context. Universal primers were incorporated into the sequence to facilitate amplification of the targeted mutation sites by PCR (**Figure 2A**). The two transgene constructs were named “TG1” and “TG2”, and together contained 35 mutation target sites to be targeted named “A-m1” through “A-m24”, “C-m1” through “C-m11”. TG1 also contained the *DNMT1* site, a control site that we previously targeted in cells and mouse retina.^45^ (**Table 1**). Please note that in this study, we use the convention that a target site named “A-m1” refers to the first *mutation locus* intended to be edited by an adenine base editor (**Table 1** “Mutation Locus” column), while the term “A1” refers to the adenine basepair in *position* 1 counting from the 5’ end of protospacer sequence (**Table 1** “Target base position” column, **Figure 1B**). The transgene TG1 or TG2 construct was inserted into the AAVS1 locus of HEK293T cells to generate stable transgene cell lines using the HDR-mediated CRISPR/Cas9 system (**Figure 2B**). Correct integration of the transgene into the *AAVS1* site was confirmed by PCR and sequencing (data not shown).

**Figure 2.**
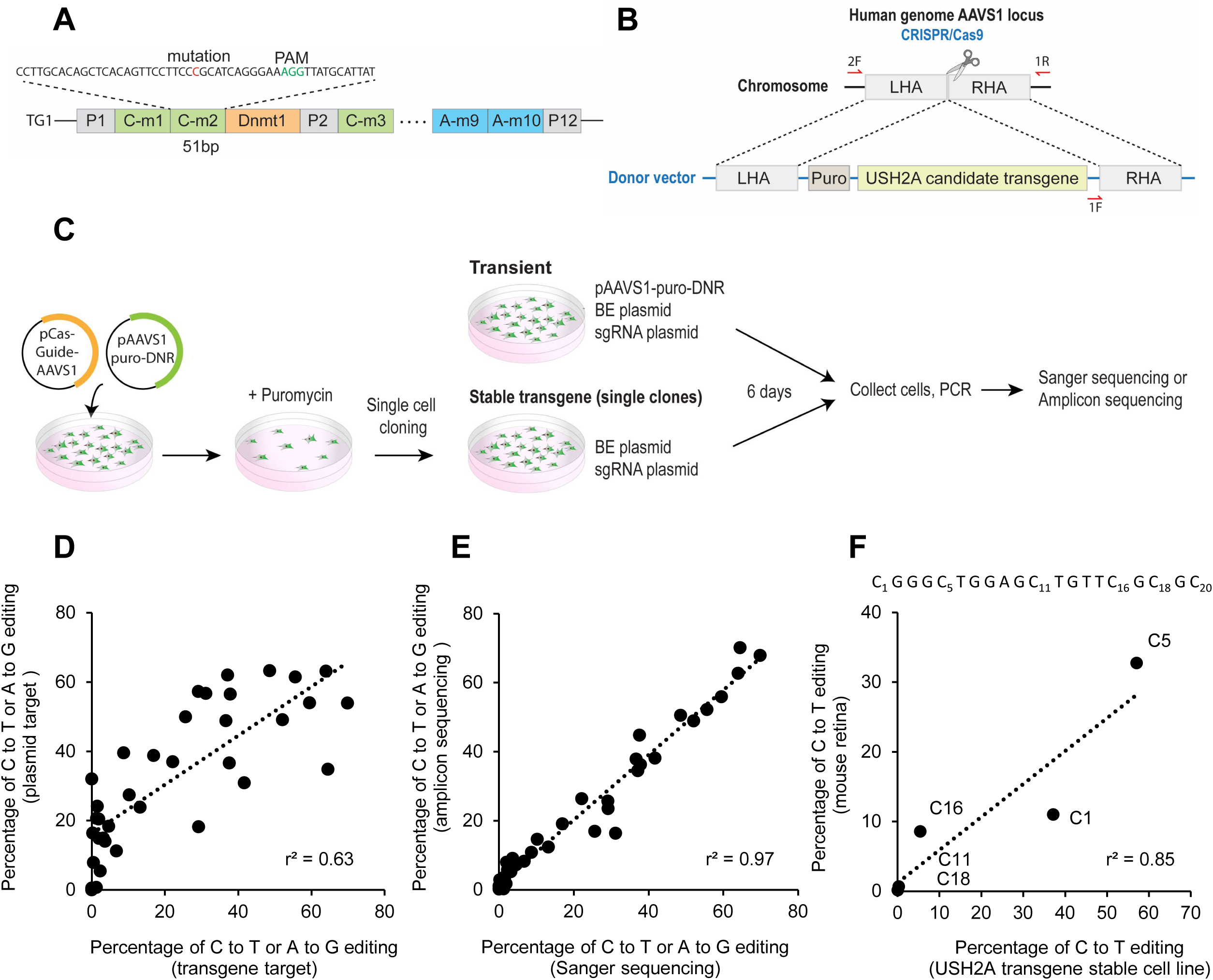
System for evaluating base editing on multiple target mutations. **(A)** Schematic showing the transgene consisting of universal primers and a tandem array of 51 bp human USH2A sequences containing pathogenic mutations. **(B)** Strategy for establishment of the transgenic, stable HEK293 cell lines, using a donor vector (p*AAVS1*-puro-DNR) targeting the *AAVS1* site by HDR **(C)** Experimental protocol for establishing the transgene stable cell line; and base editing of candidate sites in transient transfected plasmids transient and stable transgenes. **(D)** Comparison of editing efficiency of the target sites on a transiently transfected plasmid versus on a stably integrated transgene, using Sanger sequencing. **(E)** Similarity between base editing efficiency quantification using Sanger sequencing versus NGS amplicon sequencing. **(F)** Comparison of the editing efficiency in the Dnmt1 locus in the stable cell line versus mouse retina. Labels (e.g.C1) represent the position of the edited cytosine in the protospacer sequence, shown above.

### Optimizing the evaluation of base editing efficiency in vitro

We compared the base editing results between an exogenous plasmid-based target introduced by transient transfection versus a stably integrated transgene target incorporated into the *AAVS1* genomic locus (**Figure 2C**). For plasmid-based targeting, base editing reagents (a BE plasmid and a separate guide plasmid) were co-transfected with the *USH2A* transgene plasmid. For transgene-based targeting, only the base editing reagents (a BE plasmid and a separate guide plasmid) were transfected. Six days after transfection, each target sequence was amplified by PCR followed by Sanger sequencing to quantify the editing efficiency of the target base. The results showed a reasonable correlation between plasmid-based and transgene-based editing (**Figure 2D**, r^2^= 0.62).

Next, we compared two different methods of measuring editing efficiency, next-generation sequencing (NGS) versus Sanger sequencing. Sanger sequencing was optimized to have low background. PCR products of edited DNA were subjected to either NGS amplicon sequencing or Sanger sequencing, and the editing efficiency was defined as the target base conversion rate. These two data sets showed a high correlation (r^2^=0.96, **Figure 2E**), validating Sanger sequencing as an accurate method for evaluating base editing efficiency, at least when the trace background is low.

To estimate how these results derived from the HEK293 cell line might compare to editing in the retina in vivo, we compared the editing efficiency at the control *DNMT1* site in the HEK293 transgene stable cell line to our previously published editing data in mouse retina after subretinal AAV delivery of a split-intein formulation of BE3.9.^45^ The target cytosine C5 and neighboring cytosines within the protospacer sequence of *DNMT1* showed a similar editing pattern when comparing subretinal administration in vivo to results seen in cell culture (r^2^=0.85 **Figure 2F**). (Similar results for the A-m1 locus are presented below.) These results demonstrate that the editing efficiency and the editing pattern of base editing in the HEK293 cell line experiments presented in this study likely reflect those in the endogenous genome of photoreceptors in vivo, at least as indicated by results from the *DNMT1* site.

### Evaluation of base editing efficiency on *USH2A* target mutations

Due to the reasonable but imperfect correlation of editing efficiency between editing of transiently-transfected plasmids and stably integrated transgenes, the remaining experiments were performed using the stably-integrated transgene approach. Each transgene is in the autosomal genome, like the eventual therapeutic targets in photoreceptors. In each of the 11 target sites with an intended C-to-T or G-to-A target conversion (sites C-m1 to C-m11), we tested editing using one of three different PAM variants (SpCas9-WT, SpCas9-VRQR, and SaCas9-KKH) of the C-base editor CBE3.9max, using Sanger sequencing as a readout. The results showed CBE editing efficiencies ranging from 59.5 ± 8.4% (site C-m1) to 1.9 ± 1.9% (site C-m10) (**Figure 3A**). The editing window was rather wide, similar to previous reports,^41^ with editing observed of additional nearby/“bystander” cytosines in addition to the target cytosine (**Figure S2**). Next, we examined the editing efficiency of two A-base editors, ABE7.10 and ABE8e, on 24 target sites with an A-to-G or G-to-A intended target, using one of four different PAM variants (SpCas9-WT, SpCas9-VRQR, SpCas9-NG, and SaCas9-KKH) (**Figure 3B**).

**Figure 3.**
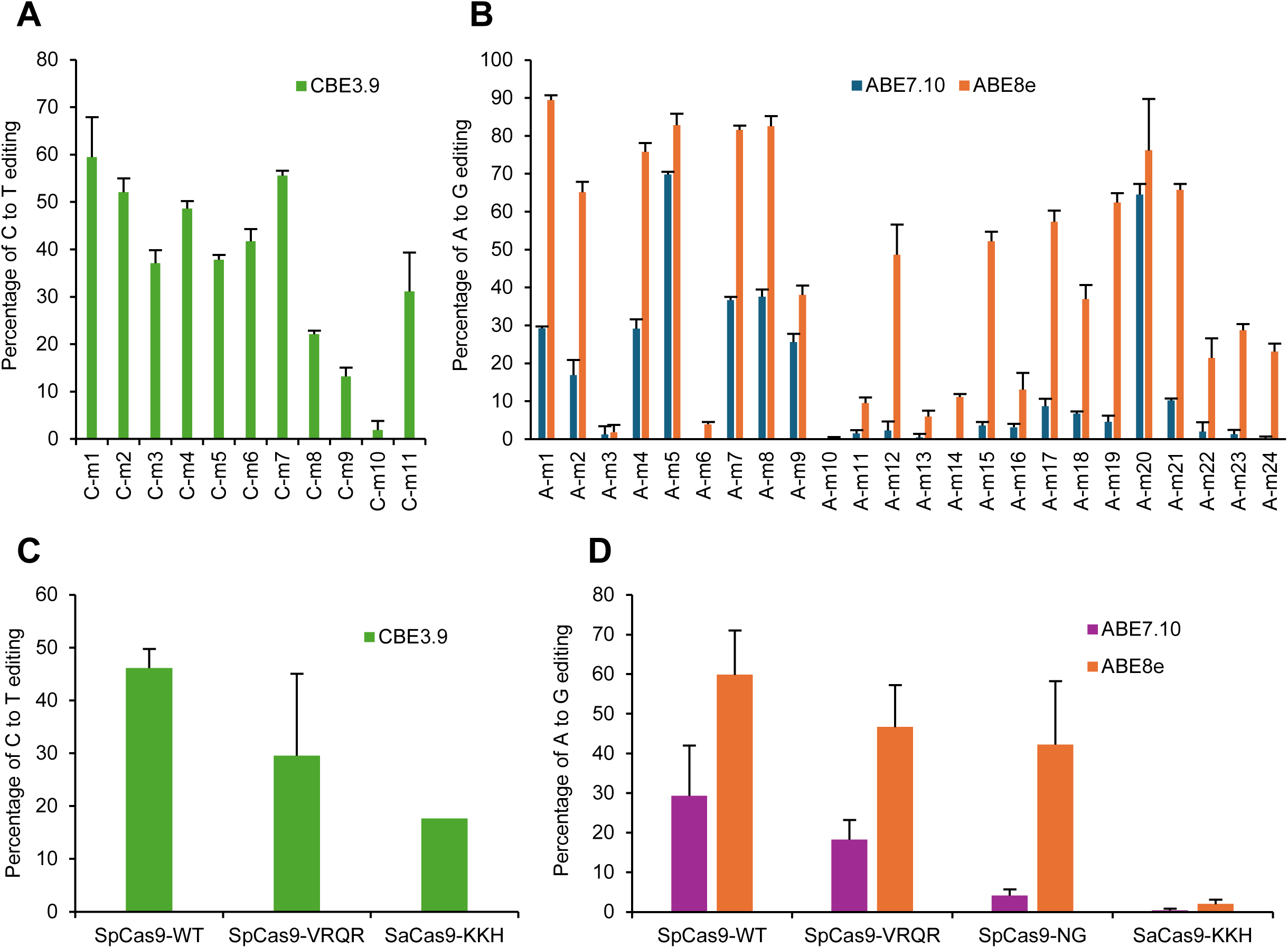
Editing efficiency of base editor/guide combinations on all target mutations, measured using Sanger sequencing with EditR quantification, in stably transfected cell lines with USH2A target mutations. **(A)** C to T editing efficiency of CBE and **(B)** A to G editing efficiency of ABE7.10 and ABE8e are shown, with mean ± SD (n = 3). **(C)** Average editing efficiency of all CBE targets, grouped by the three different Cas9 PAM variants used. **(D)** Average editing efficiency of all ABE targets, grouped by the four different Cas9 PAM variants used.

ABE7.10 showed editing efficiencies from 69.9 ± 0.7 (site A-m5) to 0% (sites A-m6, A-m10, and A-m14). ABE8e showed editing efficacies from 89.4 ± 1.3% (site A-m1) to 0.20 ± 0.35% (site A-m10). Compared with CBE, ABE tended to show greater variability as to whether a particular editing target showed efficient editing or not. As previously reported,^48^ ABE8e was more efficient and had a wider effective editing window than ABE7.10 (**Figure S3A, S3B**). Efficiency also varied by the Cas9 PAM variant type used. SpCas9WT tended to have the highest editing efficiency and SaCas9-KKH tended to have the lowest (**Figure 3C,D**).

To determine which target sites are most suitable for producing an expected phenotypic recovery at the amino acid level, the effects of bystander editing were evaluated using NGS and analysis of individual reads. “Productive edits” were defined as reads in which the intended amino acid change was created but no other amino acids were changed (**Figure 4A**). This allows for synonymous mutations but does not allow for missense mutations, nonsense mutations, or indels. This is a conservative definition in that some missense mutations and in-frame indels may be tolerated, which can be verified as needed in future work. (See Methods for “productive edits” definition for splice site mutations.) The fraction of productive edits, compared to total edits, was very low for sites C-m2, C-m3, C-m4, C-m5, and site A-m5, whereas the fraction of productive edits was high for sites C-m1, A-m1, A-m7 and A-m8 (**Figure 4B**). The indel ratio in each target was also calculated from the results of NGS amplicon sequencing, since the indel ratio is also an important parameter for evaluation of the therapeutic potential of a target editor. The indel ratio for CBE3.9, ABE7.10, and ABE8e editors ranged from 0.5-16.5%, 0-2.7%, and 0.07-1.7%, respectively (**Figure 4B**). While the overall trend in editing efficiency among editors was ABE8e > CBE3.9 > ABE7.10, the trend for indel creation was CBE3.9 > ABE7.10 > ABE8e (**Figure 4C, D**). Thus, ABE8e had the most favorable combination of editing efficiency and low production of indels. Overall, these results demonstrate that multiple *USH2A* mutations show empirical evidence of high productive edits with low indels, making them candidates for continued evaluation as therapeutic targets by base editing.

**Figure 4.**
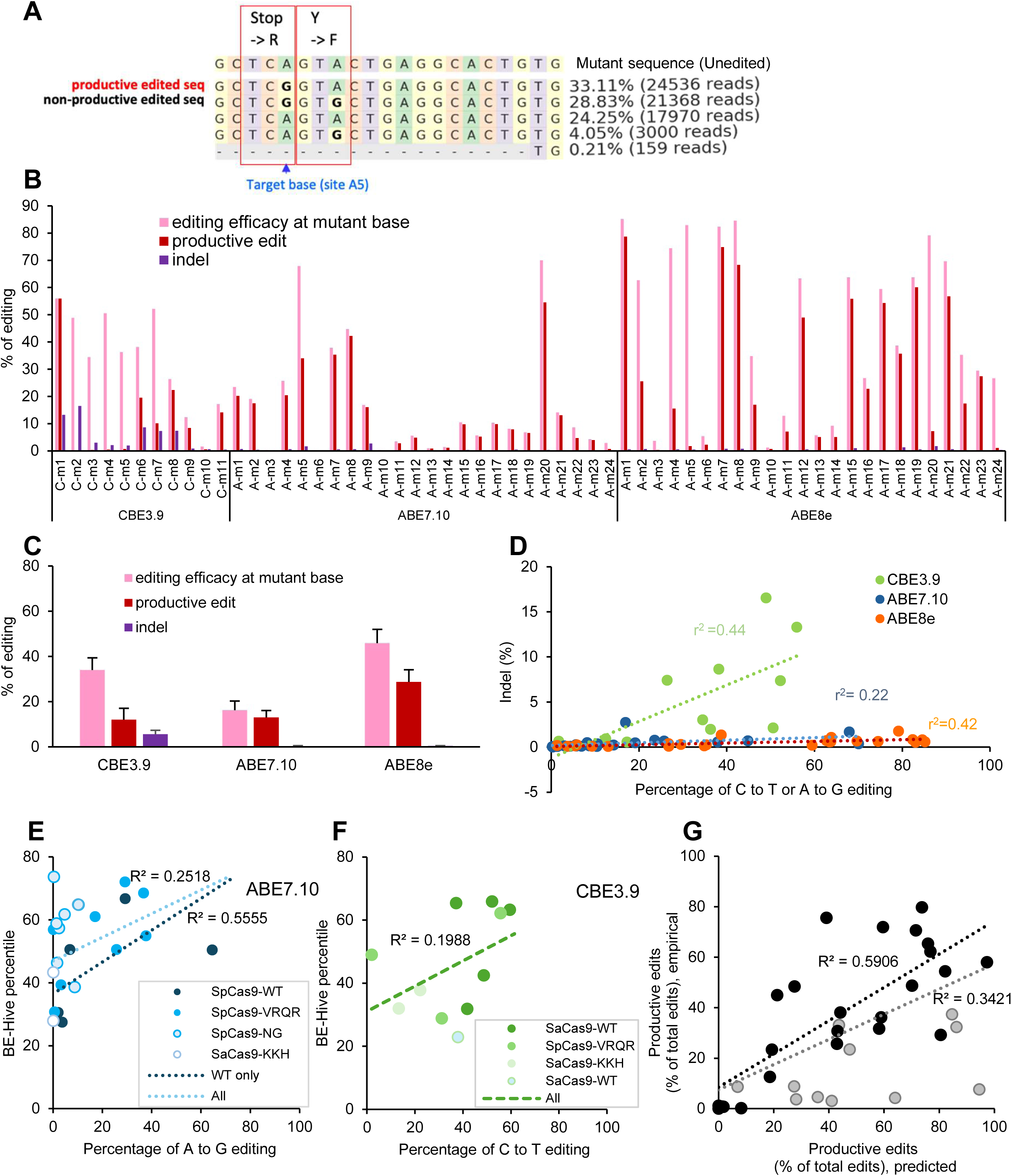
Editing efficiency of base editor/guide combinations on all target mutations, measured using NGS analysis, in stable transgene cell lines with *USH2A* target mutations. (**A**) Example editing pattern around position A5 of mutation, A-m5. A “productive edit” is defined as an edit in which the amino acid sequence is changed to wildtype. A “non-productive edit” is defined as an edit with unwanted amino acid changes. (**B**) Editing efficiency at each mutant site is shown at the mutant base itself, and as the productive editing rate summed over all observed alleles. The indel rate includes all insertions or deletions detected. (**C**) Average editing efficiency and productive editing rate is shown for the three base editors used. (**D**) Editing efficiency at mutant base vs indel frequency, with each point representing one mutant site. **(E, F)** Comparison between the experimental value of editing efficiency and predicted percentile value by BE-HIVE using ABE 7.10 **(E)** and CBE3.9 **(F)**. Point shading indicates the type of Cas9 PAM variant. The dark blue and light blue (**E**) or light green (**F**) regression lines were calculated on the data of only ABE-SpCas9-WT and all ABE SpCas9 variants, respectively. **(G)** Correlation between the *pattern* of editing (as measured by the fraction of all edits which were productive edits) as measured experimentally versus as predicted by BE-HIVE. The correlation is higher for the subset of mutant sites which have >15% editing efficiency (black dots, black regression line), compared to all sites (black and grey dots, gray regression line).

To determine the correlation between these empirical results and the editing properties predicted by BE-HIVE computational predictions^49^, we compared the empirical percent editing to the BE-HIVE percentile, which represents a predicted editing efficiency (**Figure 4E-F**). Correlations were higher for the SpCas9-WT editor compared to the PAM variant editors. We compared the editing patterns among nucleotides within each locus for both empirical and predicted results (**Figure 4G**). Specifically, the percentage of all edits that are productive edits is calculated for both the empirical and predicted results. For sites with low empirical editing efficiency (<∼15%), the pattern of editing showed less agreement between empirical and predicted results, so the correlation is calculated separately within the high- or low-editing efficiency sites (**Figure 4G**).

### Evaluation of off-target effects of base editing for *USH2A* mutations

The specificity of base editing depends on how efficiently the sgRNA will recognize other, off-target sites in the genome. Sites C-m6 and A-m1 were selected for further off-target evaluation, due to their high base editing efficiency (above). The CRISPOR *in silico* prediction algorithm^50^ was used to identify 10 sequences with a high MIT-off-target score for each target (**Table S1**). After editing HEK cells, the off-target loci were amplified by PCR (**Table S3**) and quantified by Sanger sequencing. Off-target editing rates were generally low, but the highest off-target edit with the C-m6 editing reagents was 19.4%, while the highest off-target editing rate for A-m1 was 74.5% (**Figure 5A, B**). There was little correlation between the MIT-off-target score and editing efficiency (**Figure 5C**). Because the two loci with high off-target editing efficiencies were located in an intronic region (C-m6) or in an intergenic region (A-m1), they are less likely to affect gene expression levels. More comprehensive tests for off-target editing,^51^ as well as whether any off-target edits have a deleterious effect on photoreceptor health, should be considered for future work. These results suggest that sites C-m6 and A-m1 are promising therapeutic targets for base editing.

**Figure 5.**
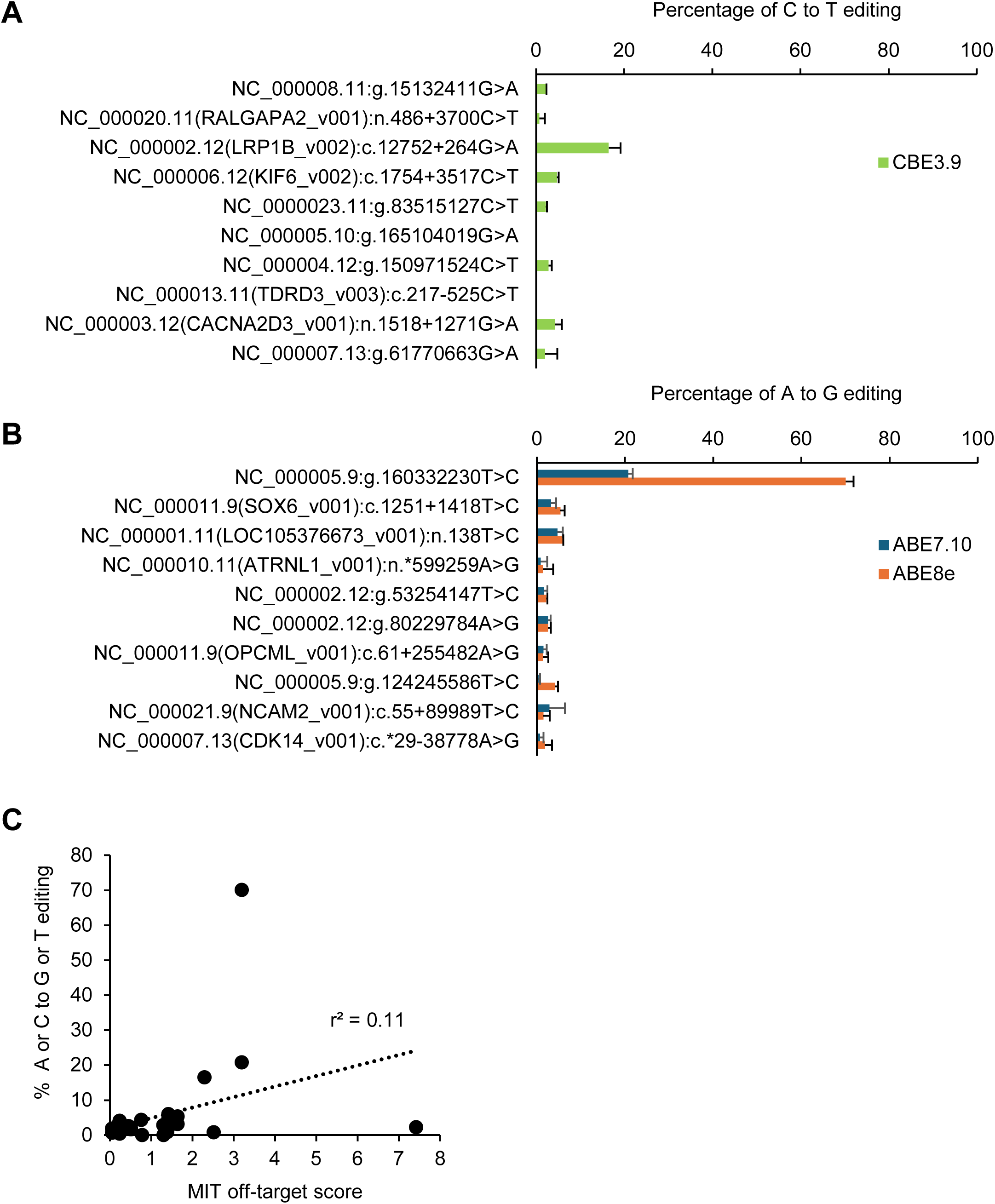
Off-target editing. The top ten predicted off-target sites for C-m6 **(A)** and A-m1 **(B)** were identified using the CRISPOR prediction tool. Editing rates are shown as mean ± SD (N=3; Sanger sequencing). (**C**) A low correlation is seen between the between MIT off-target score and the empirical editing rate at off-target sites.

### Recovery of full length USH2A protein with base editing in cells

The experiments above (**Figures 3-5**) were performed on transgenes integrated into the *AAVS1* genomic locus, using a sequence context of 51bp from each mutation locus. To test whether base editing has the same effect at the endogenous, full-length *USH2A* genomic locus, we established a mutant HEK293 cell line with a homozygous site A-m1 mutation (*USH2A* c.11864G>A p.Trp3955*)(**Figure 6A**). (Homozygous, in this case, means that all three of the triploid chromosomes in the HEK cell genome were mutagenized.) Editing of the A-m1 site in the endogenous *USH2A* locus using ABE7.10 and ABE8e showed an editing efficiency of 23 ± 1.6% and 75 ± 0.5% respectively (**Figure 6B**). These results are very similar to those obtained from editing the synthetic construct containing the A-m1 site at the *AAVS1* locus, which showed 29 ± 0.5% editing with ABE7.10 and 89 ± 1.3% editing with ABE8e. Using ABE7.10 to edit the same site in a transfected plasmid showed a lower editing efficiency of 18 ± 0.2%.

**Figure 6.**
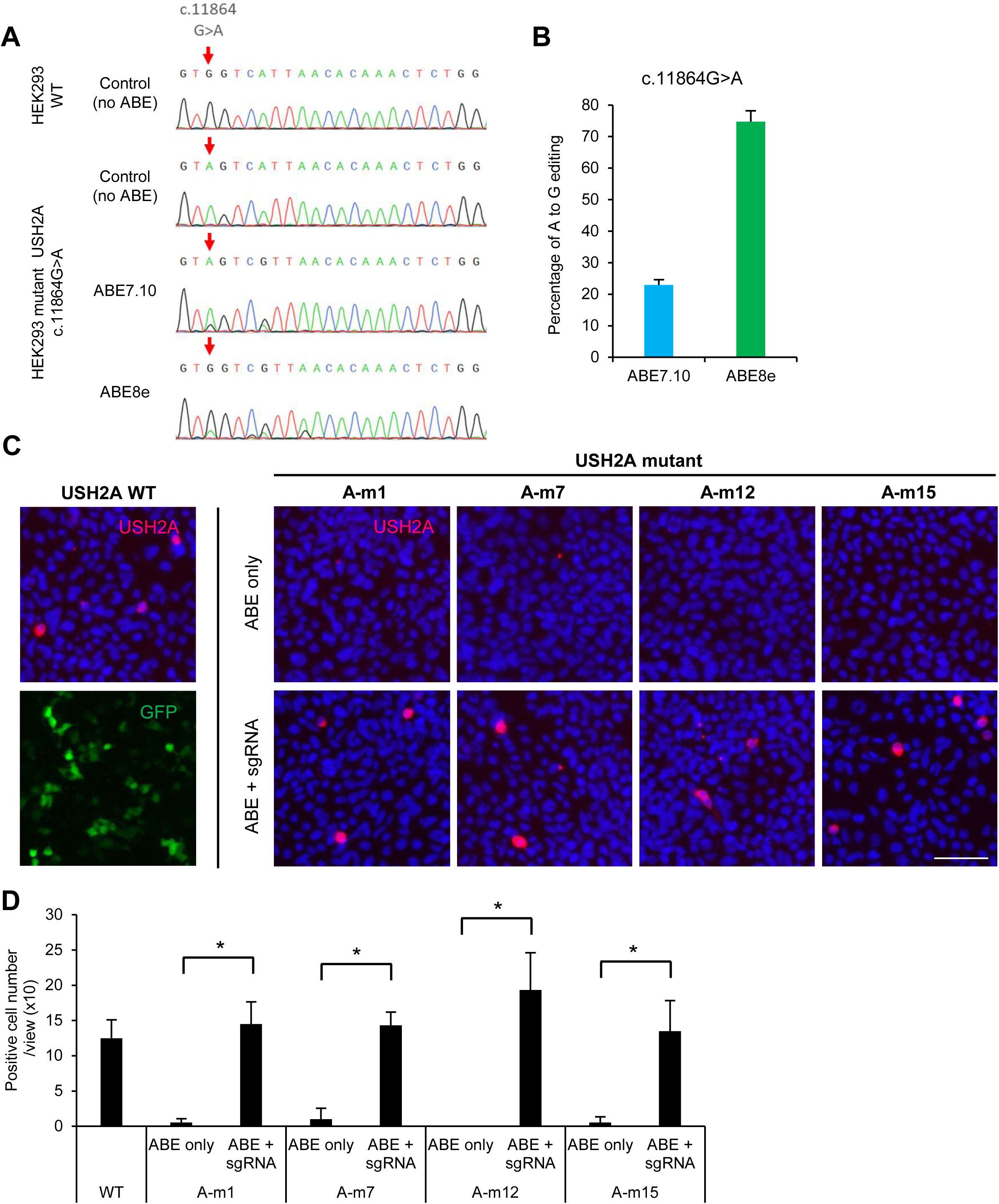
Restoration of USH2A protein expression with base editing in cells in vitro. **(A)** Sanger sequencing shows that the A-m1 mutation (c.11864 G>A; red arrow) was introduced into the native *USH2A* genomic locus, in all triploid chromosomes of a HEK293 cell line (red arrow). **(B)** This cell line was edited with ABE7.10 or ABE8e, and the editing efficiency was quantified. (**C**) Immunostaining of USH2A in HEK293 two days after transient co-transfection with plasmid vector of a full length USH2A cDNA wildtype (WT) or mutant (A-m1, A-m7, A-m12, A-m15) plasmid and base editor/ sgRNA plasmid vector. USH2A full length protein is stained (red) with an USH2A antibody that recognizes intracellular region. Nuclei are stained with Hoechst 33342 (blue). Bar; 50μm. GFP (lower left; a representative separate example) serves as a transfection control and is no longer detectable after permeabilization for USH2A staining (other panels) (D) Cells positive for USH2A full length protein were counted in 6 fields (mean ± SD, n=6), * p < 0.001, by two-tailed Student’s t-test.

Next, we examined how efficiently mutant *USH2A* protein expression is restored following base editing of nonsense mutations at A-m1, A-m7, A-m12 and A-m15, sites which were edited at high efficiency in the above experiments. We modified a wildtype *USH2A* expression plasmid^52^ by mutagenesis to produce the four individual *USH2A* mutants listed. After co-transfection of *USH2A* expression vector with ABE and sgRNA plasmids in HEK293 cells, *USH2A* expression was assessed by immunostaining using an antibody to the C-terminal portion (exon 70-72) of *USH2A*. (The modest percentage of transfected cells was attributed to the very large *USH2A* cDNA plasmid size.) In contrast, no significant immunostaining was detected from any of the four mutant proteins (**Figure 6C**). Co-transfection of base editing and sgRNA plasmids fully restored *USH2A* expression to the same level as wildtype for all four mutants tested (**Figure 6C-D**). These results demonstrate that base editing can restore wildtype *USH2A* protein expression for these mutations.

### Efficient base editing of an *USH2A* mutation in the mouse retina in vivo

Site A-m1, c.11864G>A p.(Trp3955*), was selected for further study, because the guide-editor combination for that site showed high editing efficiency, high productive edit rate, and low indel rate in the experiments above. The c.11864G>A mutation is also the most prevalent one among all 35 tested targets. A humanized knock-in mouse model was designed and generated to test base editing in the retina in the endogenous *Ush2a* locus in vivo. The G>A mutation was installed, in addition to 3 more basepair changes where the mouse DNA sequence differed from the human sequence within the protospacer region (**Figure 7B**). Creating a longer area of humanized adjacent sequence showed similar editing efficiency in vitro (not shown) and was not pursued further. Homozygous humanized mutant knock-in mice underwent subretinal injections with a pair of adeno-associated viruses (AAVs) containing a split-intein version of ABE8e^45^ and an expression cassette for the guide sequence for site A-m1 (**Figure 7A**). Virus expressing eGFP was also mixed into the injected solution as a tracer. NGS analysis confirmed the editing efficiencies at the target site were 32.3 ± 3.5% for whole retina and 64.9 ± 2.8% for sorted, transduced cells (**Figure 7C, D**). After removing reads with non-productive edits (unwanted amino acid changes), the editing efficiency was 52.3 ± 2.6% for sorted, transduced cells (**Figure 7C, D**). The pattern of adenine editing within the protospacer region was very similar between the stable cell line and the mouse retina (r^2^=0.97). Immunohistochemistry with a C-terminal antibody showed no expression in the untreated area and local restoration of USH2A antibody staining after the editing removed the upstream stop mutation (**Figure 7E**).

**Figure 7.**
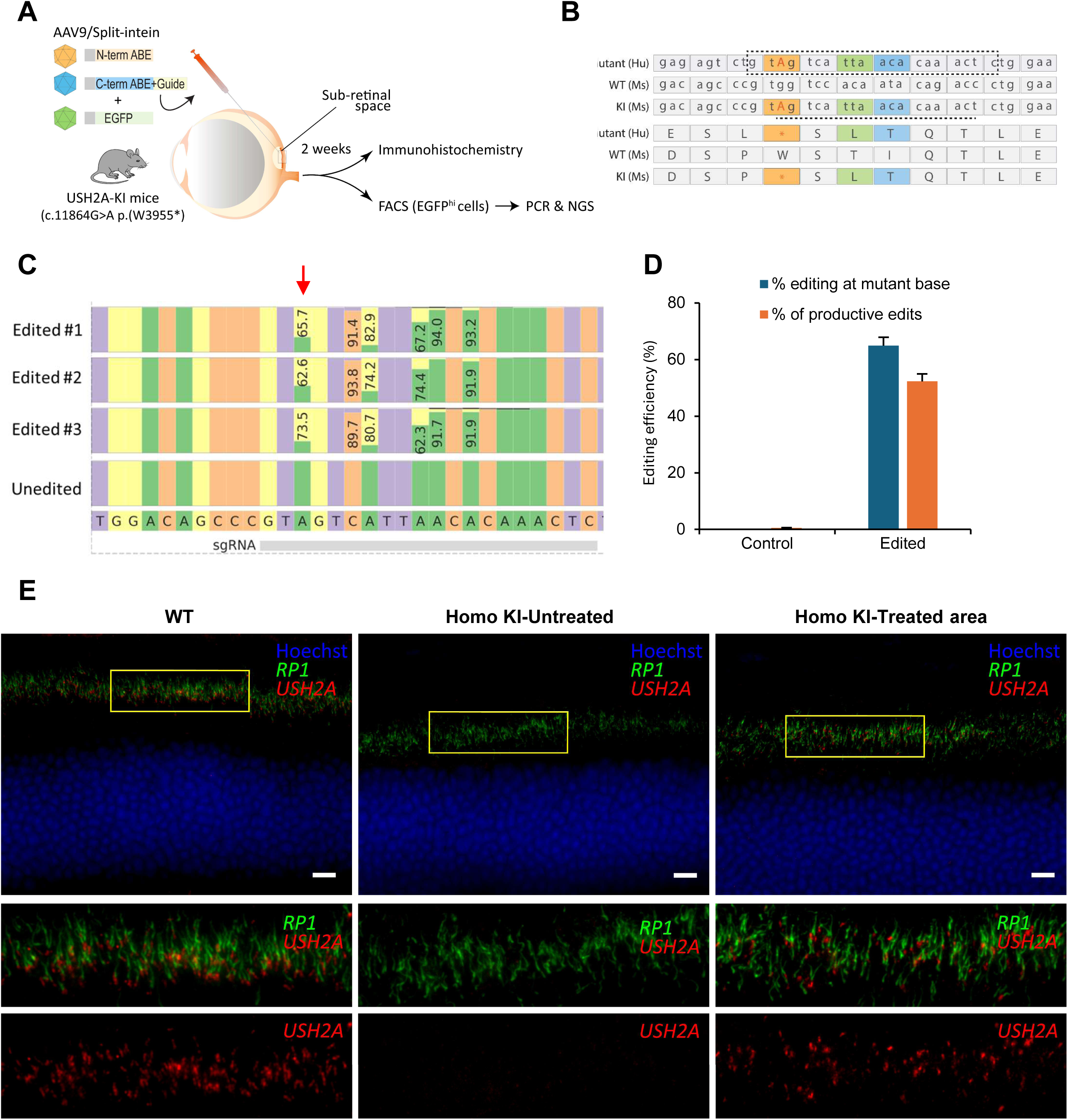
Sub-retinal delivery of ABEs via split-inteins in a humanized mouse model of the *USH2A* site A-m1 mutation. **(A)** Schema showing that 2 weeks after subretinal injection of AAV9 ABE base editors, retinas were assayed using immunohistochemistry and measurement of editing rates by next-generation sequencing (NGS) of whole bulk retina or FACS enriched EGFP^hi^ retinal cells. **(B)** The DNA and protein sequences of the human mutant, mouse wildtype, and mouse knock-in are presented, showing the guide sequence (box), the mutant site (red), and the regions that differ between human and mouse (indicated colors). Six base pairs in the underlined region within the guide sequence were humanized to generate an editable mutant sequence in the mouse genome. **(C)** CRISPResso2 analysis comparing editing efficiencies between retinas (EGFP^hi^ cells) and unedited control (N-term ABE alone + EGFP) at the target site (arrow) and bystander sites, along with the reference sequence and corresponding sgRNA binding region (bottom). **(D)** Bar graph showing the percentage of editing at the mutant base and the percentage of productive edits in the EGFP^hi^ (Cells with high EGFP expression driven by the rhodopsin kinase promoter) FACS enriched retinal cells. **(E)** Expression of USH2A in the wild-type mouse retina (left) shows USH2A antibody staining (red) at the connecting cilium and RP1 antibody staining (green) in the axoneme at the base of the photoreceptor outer segment. The outer nuclear layer is counterstained with Hoechst (blue). An untreated control eye in the homozygous knock-in (KI) retina (middle), shows no USH2A staining. In contrast, in the homozygous KI retina following base editing, restoration of USH2A staining and localization is observed (right).

Immunohistochemistry revealed the restoration of USH2A expression between the outer and inner segments of the photoreceptors (**Figure 7E**). Furthermore, the correction was further validated by co-labeling with RP1 which confirmed proper USH2A protein localization in the transition zone between inner and outer segments (**Figure 7E**)^53^. Collectively, these results indicate that ABE can be used to achieve a robust A>G transition with high efficiency in vivo. Furthermore, split-intein ABE base editors successfully restored the expression of USH2A at the protein level without requiring the delivery of the full-length cDNA sequence.

## Discussion

### Empirical validation of base editing targets

It is an exciting time in biology and medicine when the ability to revert mutations or otherwise modify particular base pairs in living cells has become increasingly feasible through techniques such as base editing. Base editing allows precise and efficient correction of SNPs without creating DNA double strand breaks and with a higher efficiency than HDR.^4,5,7^ This study provides a multi-step approach for validating base editing targets for the reversion of human mutations, using USH2A as an example. This study also demonstrated that ABE8e showed the most favorable characteristics of the base editors tested in this dataset, with high efficiency and low indels.^48^ The indel rate was lower than ABE7.10, even in the context of ABE8e’s higher editing efficiency. ABE also has a narrower editing window than CBE3.9, thus creating lesser “bystander” edits. Various approaches are now being utilized to overcome bystander activity^7,54–57^; however, many of these approaches reduce editing efficiency. While bystander editing is a potential disadvantage for mutation reversion, it is an advantage for strategies such as modifying splice sites to promote exon skipping.^58^

Different methods of empirical target validation were employed and compared, ultimately providing a helpful guide for planning similar efforts for other genes and targets. We chose to test our approach on a moderate number of USH2A mutations (N=35) that have been reported in patients. Since mutant cell lines were not available for these targets, we created and evaluated these targets on plasmids and transgenes, and the endogenous USH2A locus. Taken together, these findings indicate that for testing a single target, any of the approaches used (editing of plasmids, editing of a stable transgene fragment, editing of a mutated genomic locus) provide valid empirical data. The advantages and disadvantages of each approach are discussed below.

Sanger sequencing, which is simple and cost-effective, was found to be surprisingly accurate for quantifying the fraction of bases that are converted, compared to the gold-standard NGS-based amplicon sequencing (r^2^=0.96, **Figure 2E**). These reliable results for Sanger-based quantification were achieved in the context of low background in the Sanger sequencing traces, with background base editing estimations (i.e. of unedited bases) having a median of 0%, and very rarely above 5%. The potential decrease in accuracy when using noisy sequencing traces was not formally evaluated.

Editing of a transfected plasmid editing is the fastest and most technically convenient, in cell types that are transfectable. The target copy number may not have a large effect in this system; preliminary experiments indicated that the editing efficiency did not change significantly when the amount of the target plasmid was changed by 2-fold (not shown). Establishing a stably-integrated single-copy transgene in a particular locus (AAVS) removes the potential effect of target copy number, and was similar in complexity to creating a single mutant site in the endogenous gene, both using Cas9-based techniques with HDR templates, respectively. (Faster methods of transgenesis exist, like lentiviral transduction with antibiotic selection, the Bxb1 recombinase/landing pad system, or the piggyBac transposase system.^49,59,60^) This study used two different transgenes as additional targets were selected, but in practice, one large transgene would have been equally technically feasible. The disadvantage of editing concatenated small gene fragments into a transgene is that testing restoration of protein function is not possible, though this can be done as a separate step for top candidates (**Figure 6**). Evaluating the base editing efficiency using plasmid or target transgene-based approach is a versatile tool for studying moderate numbers of targets.

For large numbers of potential targets (hundreds or thousands), library approaches become more feasible. Innovative approaches include delivering editors and targets together using lentiviruses^59^ or using recombinase-based methods to create multiple target sites via genome engineering. Synthesizing, cloning, and deconvoluting library results is more complex and expensive than the techniques used in this study (Sanger sequencing and simple amplicon sequencing) but provide orders of magnitude more data.

Off-target experiments must be carried out in cells from the same species as the intended application. Comprehensive testing for off-target base editing is more difficult^61^ than for simple Cas9 cutting. Cas9 cutting was tested at the top 10 predicted off-target sites for the C-m6 and A-m1 guides. More comprehensive approaches (e.g. Circle Seq)^51^ should be considered for future work. It should be noted that while “perfect” guides are desired, some off-target editing may be tolerable in some settings (e.g. disease context and cell types edited), just as the off-target effects of small molecule drugs are tolerated in some settings.

### Comparison to bioinformatic predictors

As potential editing targets are selected, there are increasingly powerful methods for predicting the editing efficiency of a base editing target without experimentation, including by referencing machine learning-based models trained on large library-based experiments that provide predictions for the cell types and base editing enzymes.^49^ At some point, however, when higher confidence is needed, such as for therapeutic applications, empirical testing of individual base editing targets with actual base editing reagents is required. A collection of machine learning algorithms named BE-HIVE predict base editing efficiency and pattern.^49^ This tool was used to test the correlation between its predictions and the empirical editing data. As this work was initiated before the results of BE-HIVE models were available, it offers an opportunity to test how the model predictions held up empirically. In general, we found a moderate correlation between the model predictions and the empirical editing efficiency for the ABE editors with wildtype PAM specificities on which the model was trained. When predicting the efficiency of ABE PAM variants with lower activity, the editing efficiency was overestimated as would be expected. For CBE, the correlation was lower. Of note, only the relative, not absolute, editing efficiency is provided by BE-HIVE. The pattern of editing within the allele (i.e. target base pair versus bystander editing) is also fairly well-predicted, but only when the percentage of editing is above about 15 percent. Therefore, in situations where it is useful to detect small differences in editing efficiency and pattern between sequences, as when higher confidence is needed to optimize the best editor for an important target, such as for therapeutic applications, empirical testing of individual base editing targets with actual base editing reagents can provide the most granular data.

### Translation to an in vivo model

To validate the best-performing guide-editor combination and to compare in vitro and in vivo editing efficiencies, we generated a humanized mouse model by integrating the human mutation A-m1 (c.11864G>A, p .Trp3955*) into the endogenous mouse Ush2a locus. Retinal morphology in young Ush2a Trp3988*^ki/ki^ mice appeared normal and characterization of aged mice to evaluate for a phenotype is ongoing. Since BEs are too large to be packaged into AAV, we used split-inteins to deliver the editor and guide into the sub-retinal space. Our in vivo approach to correct an Ush2a mutation in photoreceptor cells using split-inteins gave a high percentage of productive edits (>50%) along with restoration of USH2A protein expression in treated areas. In vitro editing patterns were similar to those in the mouse retina for the Dnmt1 locus and the A-m1 locus. This outcome represents a proof-of-principle of utilizing base editors as potential therapeutic strategy for USH2A and related genes where a gene augmentation is not a feasible approach.

In conclusion, this study compared different methods of empirical evaluation of base editor-guide combinations, identifying both editors and guides with the highest efficiency. Surprisingly, a method as simple as transient transfection of plasmids followed by Sanger sequencing can provide a good empirical test of a guide-editor combination. In the process, multiple lead targets were generated for potential therapeutic base editing for Usher Syndrome type IIA, with 9 of 35 targets having productive editing rates above 50%. In particular, the A-m1 site showed efficient editing in vitro and in vivo. These experiments support the concept that base editors can broaden the addressable space of human mutations for therapeutic genome editing.

## Materials and Methods

### Ethics statement

All animal studies and examinations were performed in accordance with protocols approved by the Institutional Animal Care and Use Committee (IACUC, 2021N000068) at the Massachusetts Eye and Ear and Schepens Eye Research Institute, Mass General Brigham, Boston, and in accordance with guidelines of ARVO Statement for the use of animals and the Treaty of Helsinki.

### *In silico* selection of potentially editable human *USH2A* mutations

The *USH2A* mutation information was downloaded from LOVD (https://databases.lovd.nl/shared/genes/USH2A) and HMGD (http://www.hgmd.cf.ac.uk/ac/index.php). Likely pathogenic and pathologic mutations were selected from these databases (“UV3 as likely pathogenic”, and “UV4 as certainly pathogenic” in LOVD; “DM” in HGMD) and excluded mutations for which only 1 case was reported in LOVD. The target editing window of positions 2 to 13 (equivalent to a distance of 8 to 19 bases upstream from the 5’ end of the PAM sequence) was used to select variants that could be edited by ABE or CBE variants. The base editing PAM variants used were NGG, NGA, NG, NNGRRT, and NNNRRT.

### Plasmid vector construction

Transgenes including tandem arrays of *USH2A* mutations (**Figure S1**) synthesized by IDT were inserted into the AsCI/MluI cut site in p*AAVS1*-puro-DNR plasmid (Origene). The *AAVS1* genomic locus was selected for the transgene integration site due the availability of existing reagents targeting this locus, rather than any link to adenovirus-associated virus (AAV) biology. sgRNAs were designed to target human *USH2A* mutation (**Table S2**). Oligonucleotides were annealed and ligated into BsmBI digested BPK1520 (Addgene #65777) for SpCas9 and BPK2660 (Addgene #70709) for SaCas9 in which CAG-EGFP was inserted in HindIII cut site. The expression plasmid of the cytosine base editor, CBE3.9max,^45^ was used to make C-to-T and G-to-A transitions. PAM specificity variants for CBE editors included SpCas9-WT, SpCas9-VQR, SaCas9-WT, SaCas9-KKH, developed in the laboratory of the co-authors (DL). The following adenine base editor expression plasmids were used for mutation A-to-G to T-to-C transitions: ABE7.10 SpCas9-WT (Addgene # 102919), ABE7.10 SpCas9-VRQR (Addgene #119811), ABE7.10 SpCas9-NG (Addgene #124163), ABE7.10-SaKKH (Addgene #119815), ABE8e SpCas9-WT (Addgene #138489), ABE8e SpCas9-NG (Addgene #138491). For each mutation target site, the matching plasmids from the lists above were used, based on the editor type and PAM type.

The full-length WT human *USH2A* plasmid was driven by a CMV promoter in a pUC57 backbone.^52^ The large plasmid size (19.1kb) makes mutagenesis by PCR inefficient. Using Gibson assembly, human USH2A mutant plasmids were constructed from the WT hUSH2A plasmid. Briefly, the WT *USH2A* backbone was digested with BlpI (A-m1), SacII (A-m7), PsHAI and NotI (A-m12), or PshAI and AvrII (A-m15), which were the closest single cutters to each mutation. Two PCR fragments for each mutation were amplified from the wildtype (WT) vector with overhangs and then assembled with the cut backbone using Gibson assembly. Primers used for these PCRs are listed in **Table S4**.

### Establishment of stable cell lines containing *USH2A* mutations

HEK293 cell lines were obtained from ATCC. HEK293 cell lines containing *USH2A* mutant target sequences in *AAVS1* were established using the p*AAVS1*-Puro-DNR plasmid vector (Origene, Cat no.GE10024) and the pCas-guide plasmid vector (Origene, Cat no.GE10023). The transgenes (TG1, TG2) contained 51 bp gene fragments, each with an *USH2A* target mutation, and were connected in tandem. Universal primers were interspersed among the fragments for PCR amplification of edited sites for quantification. Synthetic oligos containing the transgene sequences were cloned into the AscI-MluI cloning site of the p*AAVS1*-Puro-DNR plasmid vector. HEK293 cells (8×10^5^cells/C6 well) were transfected with 1.6 µg of p*AAVS1*-puro-DNR and 1.6 µg pCas-guide plasmid vector using 8 µl of Lipofectamine 2000 and cultured in puromycin at final concentration of 1.5 µg/mL for 21 days after transfection. Single cells were expanded into colonies by the limiting dilution method. Insertion of the transgene in the desired *AAVS1* location was assayed using PCR with specific primers (*AAVS1*F and *AAVS1*R, **Table S2**) in the donor vector and another primer located within the *AAVS1* locus.

HDR-based CRISPR/Cas9 system was used for the establishment of a HEK293 cell line with the site A-m1 mutation (*USH2A* c.11864G>A) at the endogenous *USH2A* locus. HDR-based CRISPR/Cas9 system was used for the establishment of a HEK293 cell line with the site A-m1 mutation (*USH2A* c.11864G>A) at the endogenous *USH2A* locus. Plasmid PX459 (Addgene #48138) containing SpCas9-2A-puro and sgRNA-expressing plasmid (with guide AGGGTTCAGTGGAGAGTCTG) were co-transfected with 127bp ssODN with the c.11864G>A: TCAACAGAACTGAATGAGCACTCGTGGCTTGAGCCCAAGGAGCTGGAAAATCTTGAGGTG GAGCTTCCAGAGTTTGTGTTAATGAC**<u>T</u>**ACAGACTCTCCACTGAACCCTTGGAGTTACAGGC TCTGAC (Integrated DNA Technologies)., After puromycin selection and single-cell cloning, the sequence around the mutation site and around the edges of the homology regions was confirmed by PCR and verified by Sanger sequencing. As a result, a clone in which the mutation was contained in all 3 (triploid) chromosomes was established (**Figure 6A**). Plasmid PX459 (Addgene #48138) containing SpCas9-2A-puro was used for co-transfections with 127bp ssODN; TCAACAGAACTGAATGAGCACTCGTGGCTTGAGCCCAAGGAGCTGGAAAATCTTGAGGTG GAGCTTCCAGAGTTTGTGTTAATGAC**<u>T</u>**ACAGACTCTCCACTGAACCCTTGGAGTTACAGGC TCTGAC (Integrated DNA Technologies). After single-cell cloning, the sequence around the mutation site and around the edges of the homology regions was confirmed by PCR and verified by Sanger sequencing.

### Base editing evaluation by transfection and genomic DNA preparation

HEK293 cells (1.5× 10^5^) were seeded into 48-well plate (BD Corning) and transfected with 240 ng of base editor expression plasmid (BE3.9, ABE7.10 or ABE8) and 80 ng of sgRNA expression plasmid, and 0.66 µl of Lipofectamine 2000. The following day, the media were replaced, and the cells were maintained for six days. To obtain genomic DNA, cells were washed with PBS and lysed with DNA lysis buffer (50 mM Tris HCl pH 7.5, 0.05% SDS, and 5 µg/mL proteinase K). The genomic DNA mixture was incubated at 37 °C for 90 mins, followed by enzyme denaturation at 80°C for 30 mins.

### Sequencing and data analysis

Genomic DNA was amplified by PCR using universal primers flanking the target region. The PCR products were purified using QIAquick PCR Purification Kit (Qiagen), and Sanger sequencing and NGS amplicon sequencing was performed through service provided by CCIB DNA Core Facility at Massachusetts General Hospital, Cambridge, MA, and by the Ocular Genomics Core, Mass. Eye and Ear, Boston, MA. For Sanger sequencing, base editing efficiency was quantified by EditR (https://moriaritylab.shinyapps.io/editr_v10/). The .fastq files from NGS amplicon sequencing were analyzed by CRISPResso2 (http://crispresso.pinellolab.partners.org/). The CRISPRessoWGS Linux program was run with the following parameters: minimum homology for alignment to an amplicon 60%, quantification_window_center -10, quantification_window_size 20. The output files Nucleotide_percentage_table.txt, Alleles_frequency_table.txt, and AlleleFreqAroundSgRNATableList.txt were compiled with Microsoft Excel macros to calculate editing efficiency, indel rate, and “productive edit” rates. For missense or nonsense mutations, productive edits were defined as reads that restore the exact wildtype protein sequence, i.e. synonymous mutations were allowed, but no additional missense changes, nonsense changes, or indels were allowed. (This conservative definition may underestimate the productive edit rate if bystander missense mutations or in-frame indels are not deleterious to protein function.) For splice acceptor site mutations (C-m2, C-m3, and C-m4), productive edits were defined as reads that restored the canonical AG in the -1 and -2 positions and that also maintained the correct amino acid sequence. For the cryptic splice donor site creation mutation (C-m1), all edits which edited the C7 site (+1 splice donor site) were considered productive.

### USH2A protein recovery after base editing in vitro

HEK293 cells (1.5× 10^5^) were seeded into 48-well cell culture plates, transfected with 160 ng of ABE8 plasmid vector, 60 ng of sgRNA plasmid vector and 100 ng of *USH2A* WT or mutant (A-m1, A-m7, A-m12, A-m15) plasmid vector. Two days after transfection, cells were fixed with 1% paraformaldehyde (PFA) (Electron Microscopy Cat no. 15710) for 30 sec. After a PBS wash, the cells were blocked in 1% bovine serum albumin and 10% goat serum in PBS (pH 7.4) and were stained overnight with USH2A (1:4000) antibody that recognizes the intracellular region of *USH2A*^52^, followed by labeling with Alexa Fluor 594-conjugated secondary antibody and Hoechst 33342 for nuclear staining. Images were taken with a fluorescent microscope (Eclipse Ti, Nikon, Tokyo, Japan). Cells positive for full-length USH2A were counted in six fields per sample using ImageJ software (National Institute of Mental Health, MD, USA). Co-transfection with a GFP expression plasmid confirmed a high transfection rate. However, this GFP signal was no longer visible after permeabilization with the very mild fixation conditions described above, which are required for using the USH2A antibody.

### Mouse model for *Ush2a*

The murine endogenous *Ush2a* gene was disrupted, and human equivalent stop mutation c.11864G>A, p.(Trp3955*) was knocked into the endogenous *Ush2a* loci, exon 60 (Chr1:188874747-188874766). A schematic representation of the approach is detailed in **Figure 7B**. The microinjection was done at the Harvard Genome Modification Facility and the procedure was performed as previously described.^62,63^ Oligos and guide RNAs were synthesized by Integrated DNA Technologies (IDT Coralville, Iowa). Briefly, for targeting the mouse endogenous *Ush2a* locus, a mixture of one or two guide RNA (sgRNAs), Cas9 protein, and the HDR template were microinjected into pronuclei that were transferred into pseudo-pregnant mice. The founders were identified by Sanger sequencing (for all primer sequences, see **Table S2**). To ensure germline transmission, knock-in mice were crossed with C57BL6/J (Jackson Labs) and the resulting heterozygous mice were bred to obtain homozygous *USH2A* KI littermates. The homozygosity for *USH2A* knock-in mutation was validated by genotyping.

For genotyping, tail snips were collected and lysed using Quick Extract buffer (Lucigen Cat no.QE09050) according to manufacturer’s instructions. The target sequence of 743 bp was amplified using Promega GoTaq Master Mix (Cat no.PRM3005). The PCR product was digested with the restriction enzyme MSeI (NEB Cat no.R0525M), which resulted in two fragments (595 bp and 148 bp) for the WT and three fragments (356 bp, 239 bp and 148 bp) for the mutant. Additionally, Sanger sequencing was performed to confirm the genotype.

### In vivo base editing in the mouse retina

A split-intein construct for ABE8e and the A-m1 site guide (AAV2/9.JL961(N)-A-m1 site, AAV2/9.JL646(C)-A-m1 site), and AAV2/9.CMV.EGFP.WPRE.bGH or AAV2/9.CMV.EGFP.WPRE.bGH used as a tracer, were produced in the Gene Transfer Vector Core (GTVC) Schepens Eye Research Institute/ Massachusetts Eye & Ear.^64^ Titers were: AAV2/9.JL961(N)-A1 site, 1.17 GC/mL; AAV2/9.JL646(C) -A1 site, 3.32 GC/ml; AAV2/9.CMV.EGFP.WPRE.bGH, 5.86 GC/mL; or AAV2/9.CMV.EGFP.WPRE.bGH, 2.09 GC/ml. The sequence of gRNA is listed the **Table S1**. Sub-retinal injections were performed on mice that were 8 weeks old. Intraperitoneal injection was performed with a mixture of ketamine/xylazine (100 mg/kg/10 mg/kg) to anesthetize the mice. Subretinal injections were performed using a 33-gauge blunt needle on a Hamilton syringe and each eye received 1 µl of vectors. These vectors were administered at a high dose of 1 × 10^10^ gc/µL or a low dose of 5 × 10^9^ gc/µL.

### Immunohistochemistry

The mice were euthanized and the whole eye was dissected in PBS. To prepare retinal sections, the eyes were snap-frozen with OCT in a dry ice-ethanol slurry and stored at -80°C. The retinas were cryosectioned at 15 µm thickness onto Superfrost plus slides (Fisher Scientific Cat no. 12-550-15) using a cryostat (Leica Biosystems), air-dried, and stored at -80°C until use. The sections were post-fixed in 1% PFA for 10 min. After a PBS wash, the sections were blocked (1% BSA, in 0.1% Triton-X-100 in PBS) for 30 min at room temperature, followed by incubation with primary antibodies rabbit anti-*USH2A* (1:2000 dilution) and chicken anti-RP1 antibody (1:2000 dilution) overnight at 4^0^C. Subsequently, the slides were washed thrice with PBS, and incubated with anti-chicken Alexa flour 488 (Thermo Cat no A11039) and anti-rabbit Alexa flour 555 (Thermo Cat no. A-21428) secondary antibodies (1:1000 dilution) for 1 hour at room temperature. Hoechst (Thermo Cat no H1399) or DAPI at a dilution of 1:2000 was used for staining nuclei. After three washes with PBS, the sections were mounted onto slides using Prolong glass antifade (Thermo Cat no.P36984). Fluorescence images were captured using Nikon fluorescence microscope and the images were processed using Photoshop CS6 (Adobe) software.

### Fluorescence-activated cell sorting (FACS) of mouse retinal cells

Two weeks post injection, the mice were euthanized, and the eyes were harvested and processed as described previously.^65^ Briefly the eyes were enucleated, the lenses were removed, and the retinas were collected. Expression of EGFP in the whole retina was assessed using a Leica fluorescence microscope equipped with a FITC filter. Following a wash with PBS, retinas were digested with Solution A containing 1 mg/mL of Pronase (Sigma Aldrich), and 2 mM of EGTA (Sigma Aldrich), in BGJB medium (Thermo Cat no, 12591038) for 3 min by manual trituration to dissociate the cells. The cell suspension was subsequently treated with an equal amount of Solution B containing 100 IU/mL of DNase I (New England Biolabs), 0.5% BSA (Sigma Aldrich), and 2 mM of EGTA in BGJB medium for 5 min at room temperature. After centrifugation at 200 g for 3 min., the supernatant was drained, and the cell pellet was resuspended in 300 µl DPBS. The cell suspension was passed through a 40 µm filter prior to sorting. Sorting of EGFP-positive cells was performed on the Sony SH900 cell sorter at the Broad Flow Cytometry Core. At least 10,000 events were recorded and >25,000 EGFP^+^ cells were sorted and collected for each sample. The collected cells were subjected to DNA extraction using QuickExtract buffer (Lucigen), following manufacturer’s instructions.

### Editing efficiency analysis by next-generation sequencing

The DNA was extracted from the retina using QuickExtract buffer, following manufacturer’s instructions. PCR amplification was performed for the genomic loci of interest using Q5 Master Mix (NEB). Primers for PCR and Sanger sequencing are listed in **Supplemental Table 2**. PCR products were purified using a PCR purification kit (Qiagen) and eluted in 40 µl elution buffer (EB) and quantified with the Qubit HS DNA assay (Thermo, Invitrogen). The amplicons were then subjected to NGS. Approx ∼100 ng of purified amplicons were subjected to library preparation followed by MiSeq deep sequencing.

Purified amplicons were quantified using the Qubit dsDNA high sensitivity kit (Q33231; Thermofisher, Waltham, MA) 300-500 ng total DNA was used for NGS library preparation. NEBNext Ultra II FS DNA Library Prep kit (E7805S; New England Biolabs, Ipswich, MA) was used to ligate Illumina adapters onto DNA after enzymatic fragmentation followed by size selection for final library size of ∼320bp. Libraries were multiplexed by adding 8bp indexes (E6609S; New England Biolabs, Ipswich, MA) during the amplification step. Library quality was assessed by Tape Station 4200 HS D100 (5067-5584; Agilent, Santa Clara, CA) and quantified using Qubit HS D1000 before normalizing all libraries to 4nM concentration with a final loading concentration of 10pM after pooling. Sequencing was performed on an Illumina MiSeq instrument with v2 flow cell and a 300-cycle reagent kit (MS-102-2002; Illumina, San Diego, CA) with 2×150 paired-end reads to achieve the desired coverage. The results of amplicon sequencing were analyzed as described above.

### Data availability statement

All raw data is available upon request and the data required to evaluate the results in the paper are present within the article.

## Acknowledgments

This work was supported in part by grants from Foundation Fighting Blindness Program Project Award (QL, JC, EAP), Foundation Fighting Blindness Enhanced Career Development Award (JC), NIH NEI 1R01EY033107 (QL), NIH U01 AI142756 (DRL), NIH RM1 HG009490 (DRL), NIH R35 GM118062 (DRL) and HHMI (DRL). Core support was provided by the Ocular Genomics Institute core grant (NIH NEI P30 EY014104) and the Mass. General Hospital DNA core. The Genome Modification Facility of Harvard University in Cambridge, MA, provided transgenic service to generate humanized USH2A knock-in mice. Authors acknowledge the flow cytometry core at the Broad Institute on data collection. Authors thank Hilary Scott, Ocular Genomics Core, Mass Eye and Ear for technical assistance with Sanger and Next-Generation Sequencing. We thank Liu and Pierce lab members, Ocular Genomics Institute, Mass Eye and Ear for their critical comments, advice, and discussions.

## Author contributions

Conceptualization: Y.T., Q.L., E.A.P., and J.C.; methodology: Y.T., K.V.M., R.B., J.M.L., N.P., E.H., E.A.P., D.R.L., Q.L., and J.C.; investigation: Y.T., K.V.M., R.B., E.H., Q.L., and J.C.; visualization: Y.T., K.V.M., E.H., J.M.L., and J.C.; writing – original draft: Y.T., and J.C.; writing-review and editing: Y.T., K.V.M., J.M.L., D.R.L., Q.L., and J.C.

---

**Figure S1.**
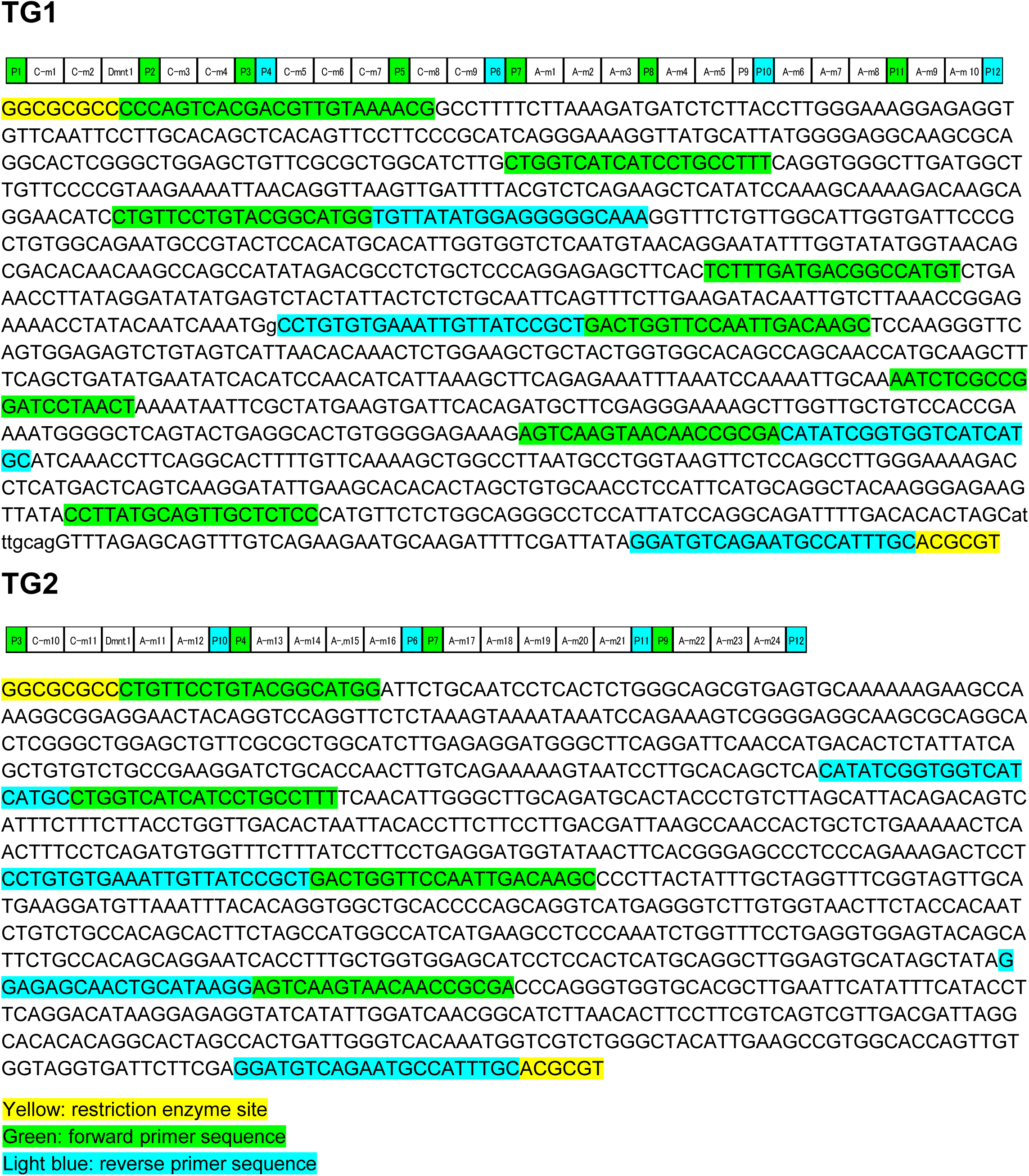
Transgene sequence with tandem USH2A mutation target sequences and universal primers (forward, green; reverse, blue) and restriction enzyme sites (yellow)

**Figure S2.**
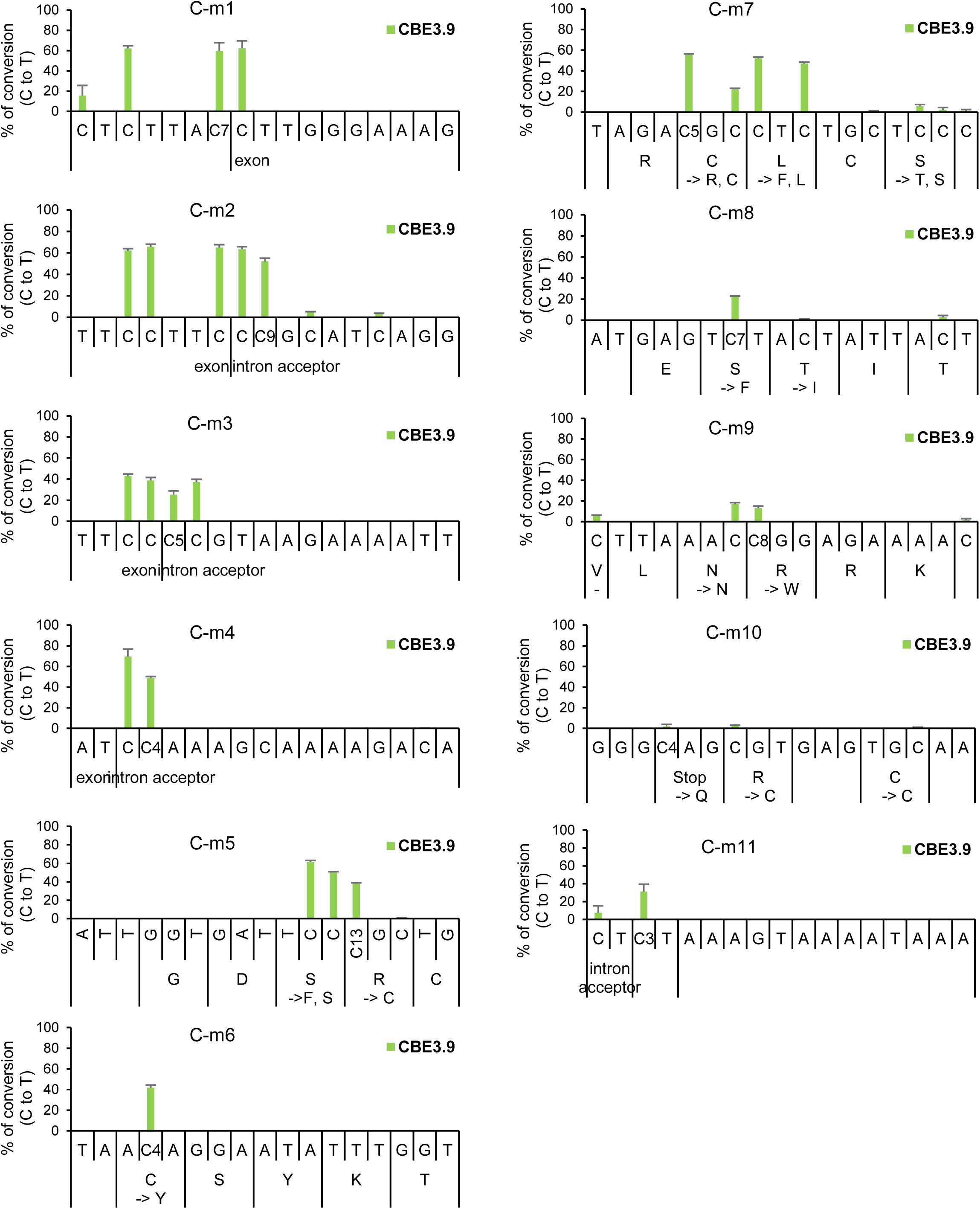
CBE editing efficiencies are shown for each basepair within the edited region. The target “C” basepair is numbered. Amino acid changes that take place with C-to-T editing are shown as indicated

**Figure S3.**
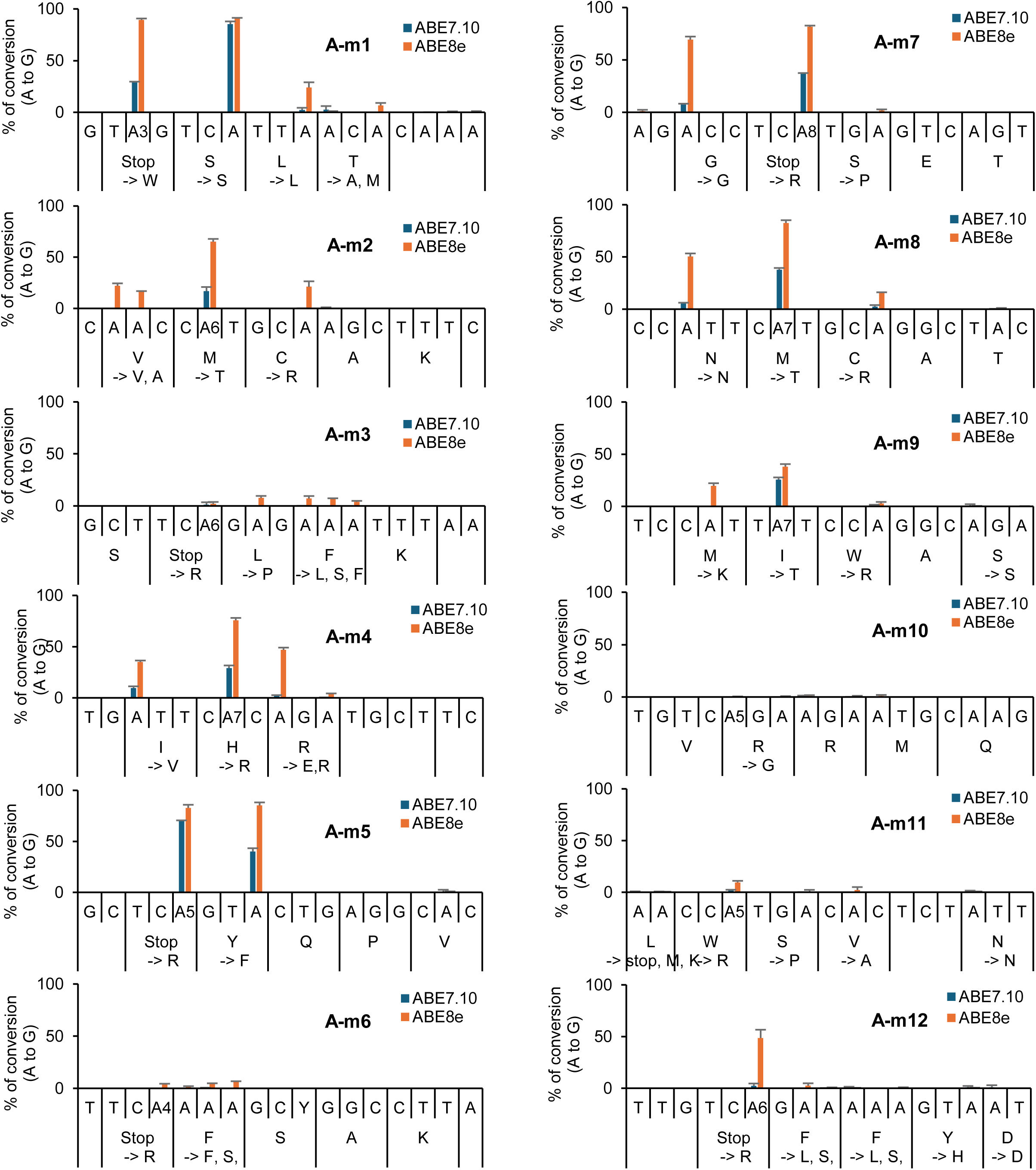
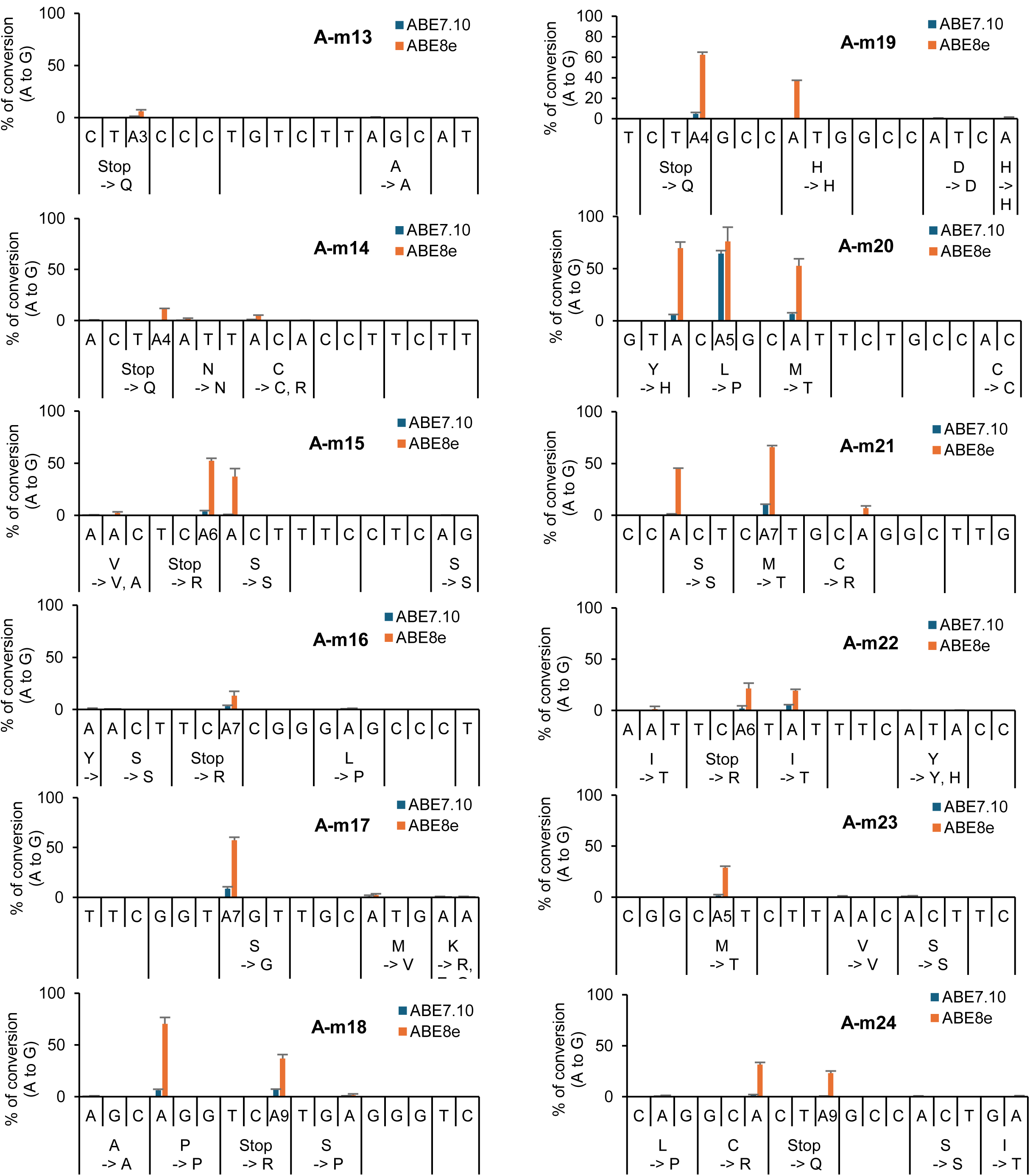
**A and S3B.** ABE editing efficiencies are shown for each basepair within the edited region. The target “A” basepair is numbered. Amino acid changes that take place with A-to-G editing are shown as indicated

**Table S1.**
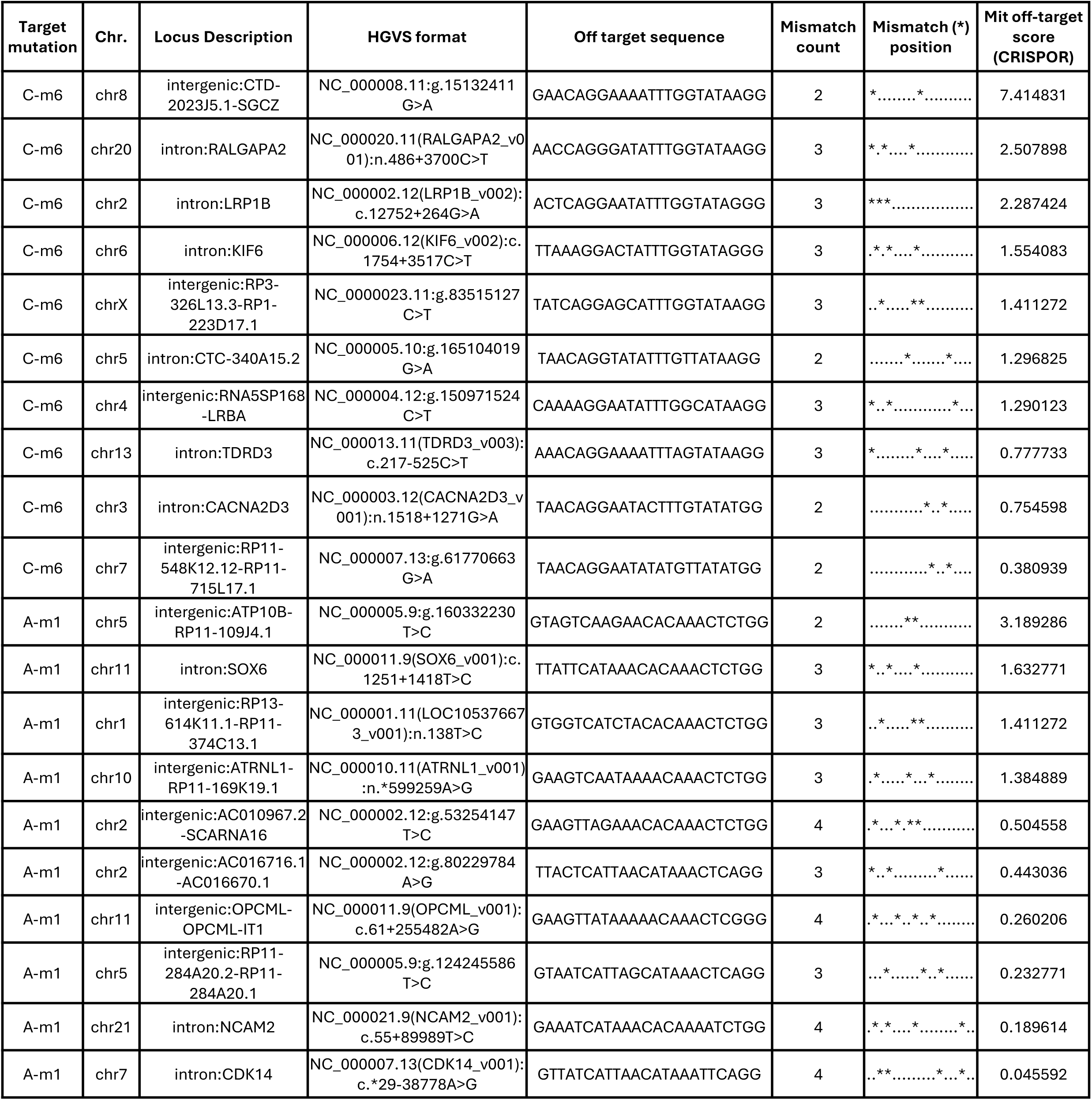
Detailed descriptions of the predicted off-target editing sites for C-m6 and A-m1, as quantified in Figure 5.

**Table S2.**
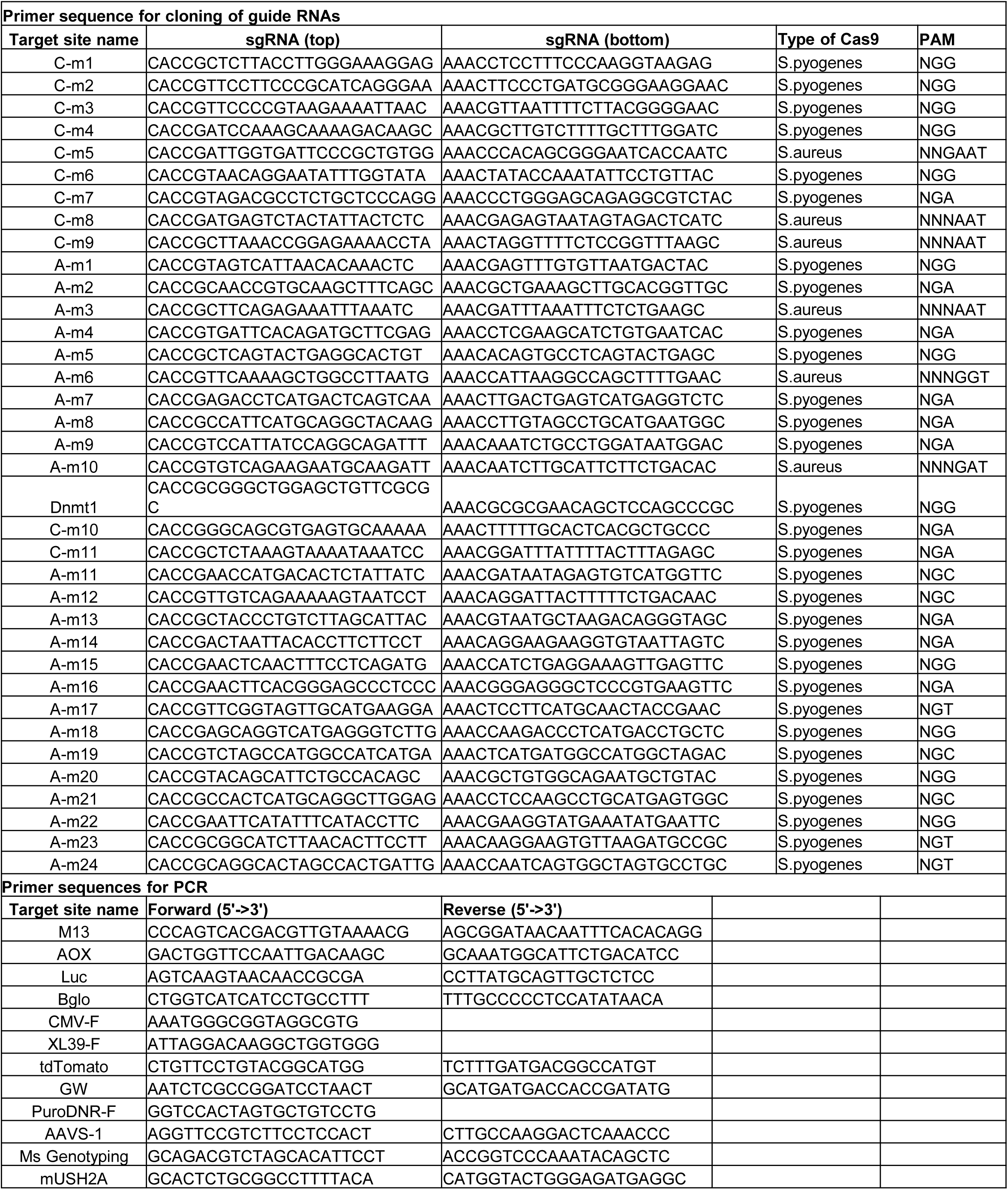
Oligonucleotide sequence for cloning of sgRNA.

**Table S3.**
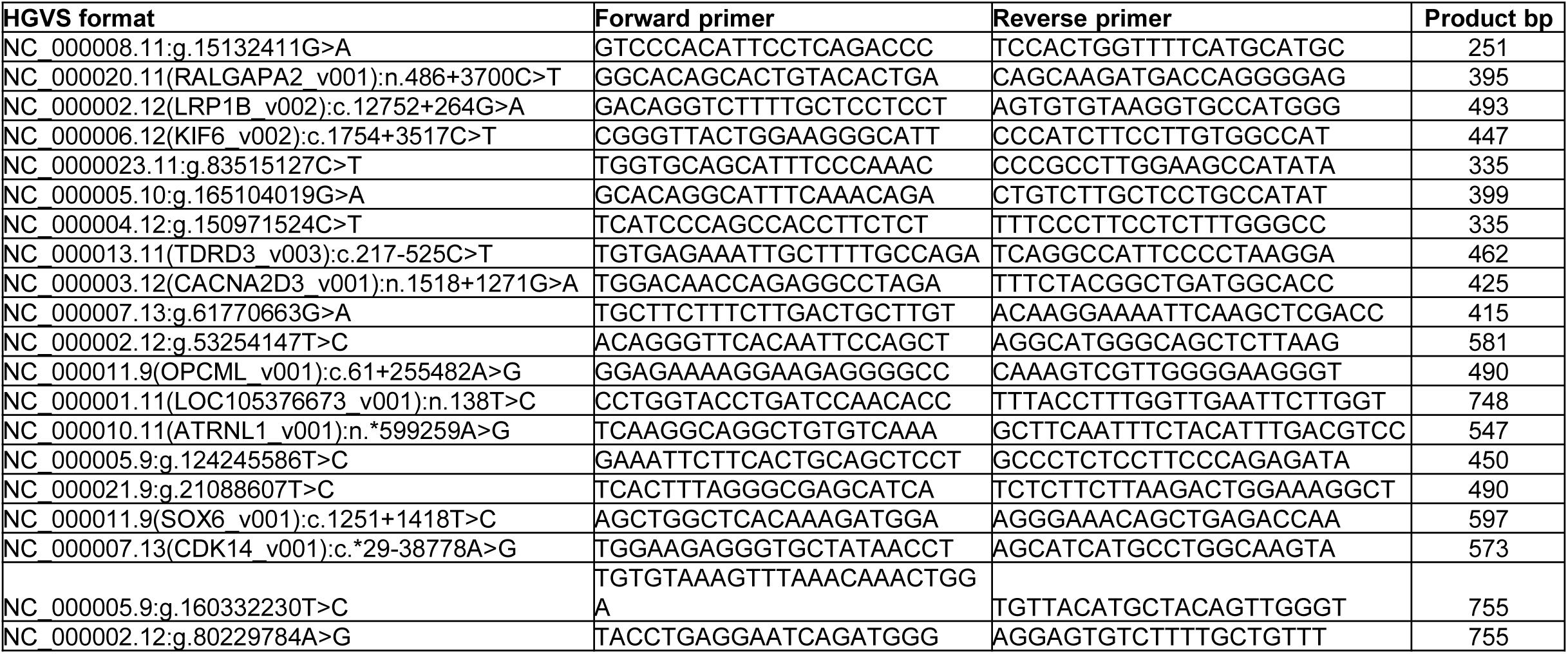
Primer sequences for off-target sequence analysis.

**Table S4.**
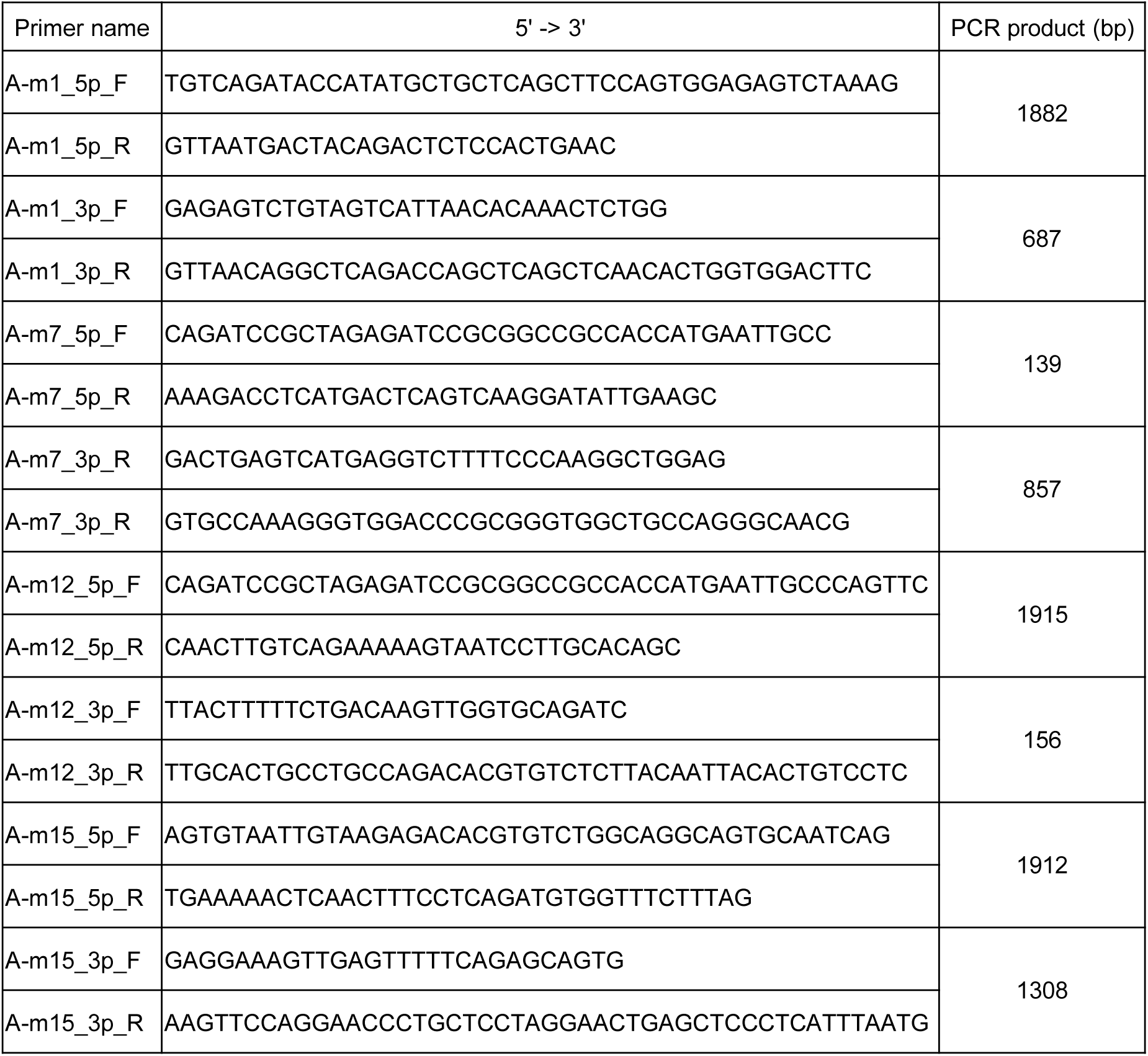
Primer sequences for cloning of USH2A mutant plasmid vector.

